# Multiplexed promoter and gene editing in wheat using the virus-based guide RNA delivery system

**DOI:** 10.1101/2022.04.06.484365

**Authors:** Wei Wang, Zitong Yu, Fei He, Guihua Bai, Harold N. Trick, Alina Akhunova, Eduard Akhunov

## Abstract

The low efficiency of genetic transformation and gene editing across diverse cultivars hinders the broad application of CRISPR technology for crop improvement. The development of virus-based methods of CRISPR-Cas system delivery into the plant cells holds great promise to overcome these limitations. Here, we applied the barley stripe mosaic virus (BSMV) for delivering guide RNAs (sgRNA) into the Cas9-expressing wheat lines to create targeted deletions in the promoter of a transcription factor and to perform multiplexed editing of agronomic genes. We demonstrate that pooled BSMV-sgRNAs could be used to generate heritable targeted deletions and multiple mutations in the genome. We transferred the high-expressing allele of Cas9 into adapted spring and winter cultivars and successfully performed the BSMV-sgRNA-based editing of two agronomic genes. The strategies presented in our study could be applied to any cultivar for creating new *cis*-regulatory diversity or targeting multiple genes in biological pathways or QTL regions, opening possibilities for the effective engineering of crop genomes and accelerating gene discovery and trait improvement efforts.

## Introduction

Crop improvement using the CRISPR-Cas-based editing relies on understanding the function of genes involved in the regulation of biological processes affecting phenotypic variation. While major advances were made towards linking genes with major agronomic phenotypes in wheat and other crops, and genome sequences facilitated inter-species extrapolation of functional information among related species, the mechanistic basis of most of the traits remains poorly characterized. The advances in gene mapping and large-scale genomic analyses helped to identify a number of quantitative trait loci (QTL) and biological pathways associated with major traits (He *et al*. 2022). However, the number of candidate causal genes detected in these studies still remained beyond the technical capabilities of existing genomic screens available for gene function validation in major crop species. The ability of CRISPR technology to introduce targeted changes into genomes has been broadly utilized in model systems to develop high-throughput functional screens greatly accelerating the characterization of causal genes and pathways (Shalem *et al*. 2015). Although, the large-scale CRISPR-Cas editing is a powerful gene discovery tool, the scope of its application in crop genetics is limited by the complexity and cost of such projects (Liu *et al*. 2020) and warrants the development of more effective gene editing strategies.

In many crops, including wheat, the realization of the full potential of CRISPR technology is hindered by a combination of methodological challenges. While CRISPR technologies based on the Cas9 and Cas12a editors have been successfully applied to edit single and multiple genes in the wheat genome(Wang *et al*. 2014; Gil-Humanes *et al*. 2017; Liang *et al*. 2017; Sánchez-León *et al*. 2018; Zhang *et al*. 2019), they show relatively low editing efficiency and required transformation of a large number of plants or screening of large populations in the next generation of transgenic plants to recover desired mutations (Wang *et al*. 2018a, 2021). Moreover, the genetic transformation protocols developed for wheat, as well as for many other crops, are restricted to few varieties showing high regenerative capabilities (Debernardi *et al*. 2020). This reduces the utility of CRISPR technology for the high-throughput editing of a large number of genes or the direct editing of adapted cultivars for testing the effects of novel CRISPR-induced alleles in diverse genetic backgrounds. The recent discovery of growth regulators *Baby boom, Wuschel* (Lowe *et al*. 2016) and *GRF-GIF* (Debernardi *et al*. 2020) significantly improved the regeneration efficiency and broadened the range of wheat cultivars amenable to genetic transformation. However, plant transformation remains a time- and resource-consuming effort that requires specialized expertise, limiting its application in most of the breeding or research programs.

Recently, virus-based CRISPR delivery systems were developed and tested for several major crops, including wheat (Čermák *et al*. 2015; Gil-Humanes *et al*. 2017; Hu *et al*. 2019; Ellison *et al*. 2020; Li *et al*. 2021). Compared to the Agrobacterium- or biolistic-based delivery of editing reagents, these systems rely on the natural ability of viruses to spread across the plant cells and allow for omitting the plant transformation and regeneration steps (Baltes *et al*. 2014). A DNA-based viral replicon was successfully used for delivering CRISPR, Cas9 and DNA templates for homology-directed repair into tomato plants (Čermák *et al*. 2015). The RNA viruses used for delivering the guide RNAs could maintain the high editing efficiency in the wheat and tobacco lines expressing Cas9 enzymes (Ellison *et al*. 2020; Li *et al*. 2021). The gene editing system based on the barley stripe mosaic virus (BSMV) was capable of generating both somatic and heritable mutations in single or multiple genes in wheat plants expressing Cas9 (Hu *et al*. 2019; Li *et al*. 2021). Because of the ease of viral infection procedure, the virus-based guide RNA delivery system could potentially be adopted by any genetic research and breeding programs and scaled up to edit many targets, overcoming the limitations of genetic transformation in wheat and other crops.

Here, we investigated the ability of the BSMV-based viral system to induce multiple targeted mutations and fragment deletions in the genomes of wheat cultivars carrying the introgression of a high-expressing allele of the Cas9 gene. By multiplexing BSMV-gRNA constructs, we created the series of deletions in the promoter of a transcription factor controlling domestication traits in wheat (Simons *et al*. 2006). According to recent studies, CRISPR/Cas9-induced *cis*-regulatory mutations in genes controlling productivity traits could broaden range of phenotypic variation and help to overcome the negative impact of epistasis on major agronomic traits (Rodríguez-Leal *et al*. 2017; Soyk *et al*. 2017). However, application of this strategy for mining beneficial *cis*-regulatory diversity remained limited in crops mostly due to the difficulties associated with production of *cis*-regulatory mutants. In our study, we demonstrate that BSMV-based viral system is an effective tool for creating new allelic variants in the regulatory regions of the wheat genome. In addition, using wheat lines with the low-and high-expressing Cas9 loci, we investigated the relationship between 1) the levels of Cas9 expression and the efficiency of BSMV-based genome editing, 2) the frequency of somatic editing and the heritability of BSMV-Cas9-induced mutations, and 3) the levels of BSMV-gRNA construct multiplexing and the efficiency of target editing. Our study shows that marker-assisted introgression of a high-expressing allele of Cas9 provides efficient means for BSMV-sgRNA-based genome editing in any adapted cultivar and opens new possibilities for the analysis and discovery of new allelic diversity for crop improvement.

## Results

### Efficiency of BSMV-based CRISPR editing depends on the Cas9 expression levels

To assess the ability of BSMV to deliver sgRNAs into wheat cells and induce mutations, we used a highly efficient QT1 gRNA (Wang *et al*. 2016) to target the coding regions of the three homoeologous copies of the *Q* gene (*TraesCS5A02G473800, TraesCS5B02G486900, TraesCS5D02G486600*), which controls major domestication traits in wheat (Simons *et al*. 2006). The Cas9-induced mutations at the QT1 site are expected to generate the loss-of-function *q* alleles, which should result in speltoid spear-shaped spikes characteristic of the wild relatives of domesticated wheat (Simons *et al*. 2006). The QT1-sgRNA was subcloned into pBSMVγPDS (Figure 1a) to replace the *TaPDS* gene fragment (henceforth BSMV-QT1) (Figure 1b). The 2^nd^ leaf of the two-leaf seedlings from the progeny of a transgenic Bobwhite line (7438) constitutively expressing Cas9 (Wang *et al*. 2016) was inoculated with BSMV-QT1 (Figure 1c). The inoculated plants showed a clear symptom of viral infection (Figure 1d). The next-generation sequencing (NGS) of the QT1 target site amplicons generated using DNA isolated from the 4^th^ leaf of plants two weeks after inoculation (Figure 1e) was able to detect the Cas9-induced mutations at the frequency of 0.43% (Figure 1f and Table S1). The target site mutations were not detected in the next generation of these BSMV-QT1-inoculated plants. We concluded that the low expression level of Cas9 in transgenic line 7438 was one of the contributing factors to such low editing efficiency (Figure S1).

**Figure 1.**
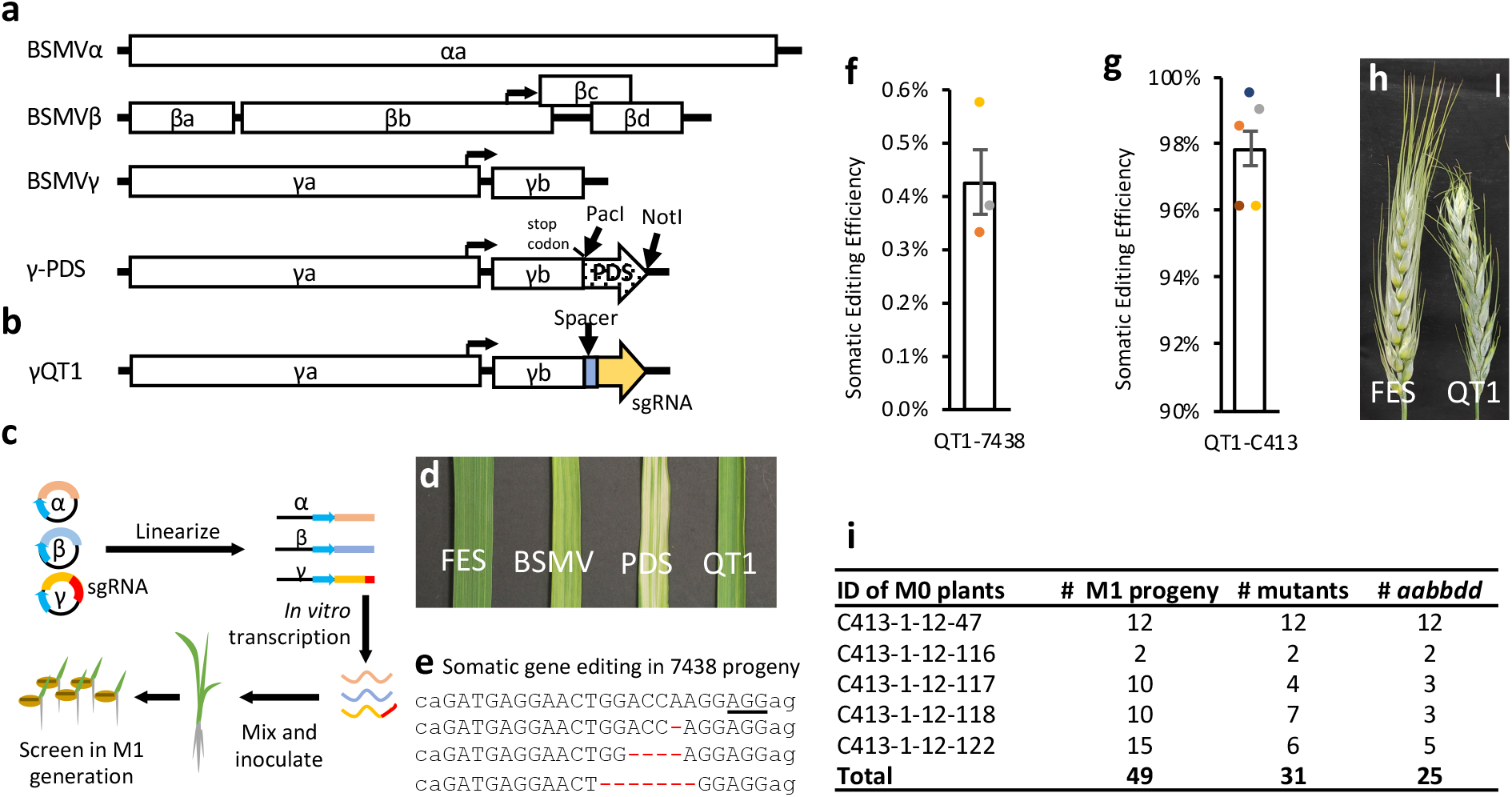
*Q* gene editing using the BSMV-based sgRNA delivery system. **a**. Structure of BSMV and control constructs used for PDS silencing. The open reading frames (ORFs), promoters and untranslated regions are shown as open boxes, arrows, and hatched boxes, respectively. Orientation of ORFs is depicted by arrow-shaped boxes. **b**. Structure of the gamma chain with QT1 guide targeting the Q gene. **c**. BSMV-gRNA-based genome editing workflow. **d**. The images of the 4^th^ leaf inoculated with the FES buffer, BSMV, BSMV-PDS, and BSMV-QT1. The 2^nd^ leaf was used for inoculation. FES buffer is the mock inoculation control; BSMV is the virus without insertion; PDS is BSMV carrying the *PDS* gene fragment; BSMV-QT1 is the BSMV-sgRNA construct targeting the *Q* gene. **e**. Examples of somatic mutations found in M_0_ plants inoculated with BSMV-QT1. The AGG PAM sequence is underlined **f**. The somatic editing efficiency in 7438-QT1 plant with low Cas9 expression (see Supplementary Fig. 1). **g**. The somatic editing efficiency in C413-QT1 plant with high Cas9 expression (see Suppl. Fig. 1). **h**. Spike morphology of the wild-type and mutant plants. The plant showing wild-type spike morphology was inoculated with FES buffer, whereas plant with speltoid spikes was inoculated with the BSMV-QT1 construct. **i**. The frequency and genotypes of Q gene mutants induced by inoculating the C413 line with BSMV-QT1.

Measuring the relative levels of Cas9 expression in multiple transgenic Bobwhite lines, we identified lines C413 and 707 (henceforth, high-Cas9 lines), both expressing Cas9 at the levels ∼ 15-fold higher than its in line 7438. Five plants derived from the high-Cas9 line C413 were inoculated with BSMV-QT1. The high levels of somatic editing efficiency, reaching 98% on average (Figure 1g and Table S1), were detected at the QT1 site in all plants. The recovery of the non-functional *q* alleles in these M_0_ plants is also supported by the speltoid spike phenotype (Figure 1h). The NGS of the M_1_ generation plants confirmed the heritability of the BSMV-induced mutations (Figure 1i). Among 49 analyzed M_1_ plants, 31 carried mutations in at least one genome copy and 25 plants were homozygous or heterozygous for mutant alleles in all three homoeologous copies of the *Q* gene (Figure 1i). This result suggests that the high Cas9 expression is required to support effective genome editing using sgRNAs delivered via BSMV.

### The efficiency of somatic editing correlates with the heritable mutation rate

To evaluate the effect of mobile RNAs on editing efficiency, Arabidopsis *Flowering Locus T* (*AtFT*), wheat *Vrn3* (Yan *et al*. 2006) and methionine and isoleucine tRNAs (tRNA^met^ and tRNA^ile^) were fused with the sgRNA targeting the *TaGW2* gene (Figure 2a). Compared to non-fused BSMV-GW2T2 construct, which showed 77% editing efficiency in inoculated leaves (Figure 2b), *AtFT, Vrn3* and tRNA^ile^ fusions, except tRNA^met^, had substantially reduced somatic editing efficiency (29%, 33% and 6%, respectively). The tRNA^met^ fusion resulted in a minor efficiency reduction reaching 76% compared to non-fused BSMV-GW2T2. Consistent with these results, gene editing efficiency reduction caused by mobile RNA fusion was also observed for a sgRNA targeting the *PDS* gene in the wheat study (Li *et al*. 2021).

**Figure 2.**
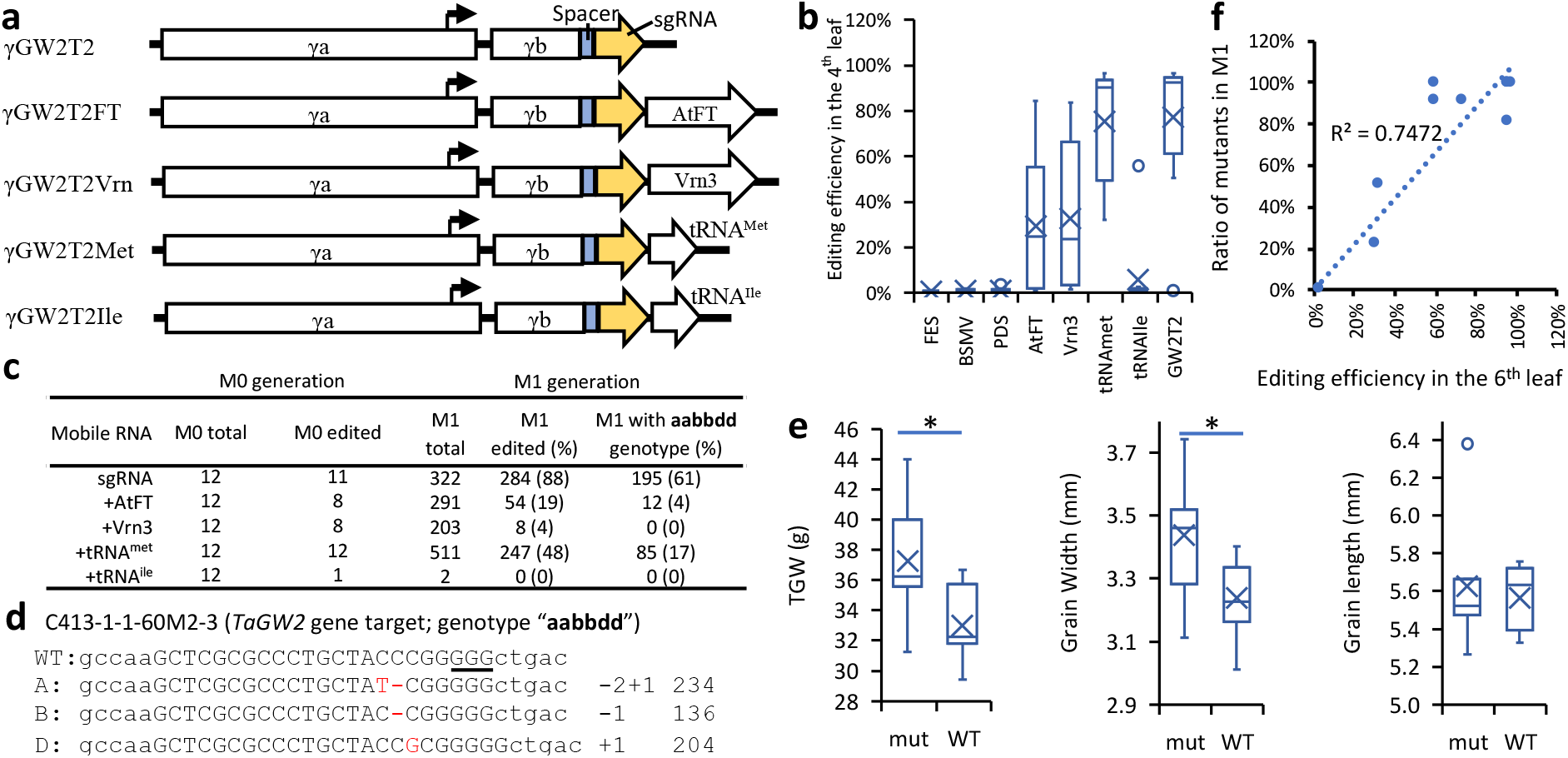
The effects of mobile RNA fusion on gene editing efficiency induced by the BSMV-sgRNA constructs. **a)** The structure of the BSMV-GW2T2-based constructs targeting the *TaGW2* gene. The GW2T2-sgRNA was fused with the *AtFT* (γGW2T2FT), *Vrn3* (γGW2T2Vrn), tRNA^met^ (γGW2T2Met) and tRNA^ile^ (γGW2T2Ile). The open reading frames (ORFs), promoters and untranslated regions are shown as open boxes, arrows, and hatched boxes, respectively. Orientation of ORFs is depicted by arrow-shaped boxes. **b)** Somatic editing induced by the BSMV-sgRNA constructs with and without mobile RNA in the non-inoculated 4^th^ leaf of the C413 line. The mutation frequency at the GW2T2 target was calculated by combining data from all three genomes. **c)** The summary of BSMV-sgRNA inoculation experiment showing the number of inoculated plants and the frequency of mutations in *TaGW2*. The somatic mutations in the M_0_ generation were evaluated using DNA extracted from the non-inoculated 4^th^ leaf. The number mutants in the M_1_ generation corresponds to the number of plants carrying mutations in at least one homoeolog of *TaGW2*. The genotypes of triple *TaGW2* mutants with mutations in each of the homoeologous wheat genomes A, B and D are shown as lower-case letters *aabbdd*. **d)** The alignments of representative NGS reads from M_1_ plant C413-1-1-60M2-3, which carries mutations in each of the homoeologous copies of *TaGW2* from three wheat genomes. The level of divergence between the wheat genomes allows for homoelog-specific alignment of NGS reads. The deleted and inserted nucleotides are shown in red. The number of deleted (−1 or -2) or inserted (+1) nucleotides are shown on the right of each read type. The number of wild-type (WT) and mutated reads aligned to each genome is shown on the right side of mutation types. The PAM sequence is underlined. **e)** Phenotypic effects of BSMV-sgRNA induced mutations in *TaGW2*. The thousand grain weight (TGW), Grain Width, Grain Length of triple mutants (*aabbdd*, n = 7) and wild type plants (*AABBDD*, n = 7) are compared. \**P* < 0.05 based on the Student’s *t*-test. **f)** Relationship between the proportion of *TaGW2* mutants in the M_1_ generation and somatic editing efficiency evaluated in the 6^th^ leaf at the booting stage of M_0_ plants. Each data point stands for an individual plant.

By genotyping the M_1_ generation, we show that the proportion of M_1_ plants with inherited target site mutations correlated with the frequency of somatic mutations detected in the leaves of the M_0_ plants. The BSMV-GW2T2 produced the highest number of mutants in M_1_ generation (88%), followed by BSMV-GW2T2-tRNA^met^ (48%) and BSMV-GW2T2-AtFT (19%). The proportion of M_1_ mutants created using BSMV-GW2T2 with all six copies of *TaGW2* mutated (Figure 2d) reached 61%. Consistent with our previous report (Wang *et al*. 2018b), the *TaGW2* mutants had significantly increased grain weight (13%) and width (6%) compared to wild-type plants (Figure 2e). We assessed the frequency of somatic mutations induced by BSMV-GW2T2 and BSMV-GW2T2-tRNA^met^ using DNA isolated from the 2^nd^ (inoculated leaf), 4^th^ and 6^th^ leaf. The frequency of somatic editing positively correlated with the frequency of M_1_ mutants, with the somatic editing in the 6^th^ leaf showing the highest correlation with the frequency of M_1_ mutants (Figure 2f and S2).

### Application of pooled BSMV-sgRNAs for editing multiple genes and generating promoter deletions

Previously, it was demonstrated that the BSMV-based system could be used to simultaneously edit multiple loci in wheat (Li *et al*. 2021). However, it remained unclear whether the efficiency of editing is affected by the BSMV-sgRNA pooling compared to the individual BSMV-sgRNA constructs. We performed multiplex editing by inoculating the high-Cas9 lines with the pool of RNAs synthesized from the BSMV-sgRNA constructs targeting *TaGW2, TaUPL3* and *TaGW7* (henceforth, the BSMV-GUG pool) (Figure 3a). Compared to plants inoculated with the BSMV-sgRNAs targeting single gene, the BSMV-GUG pool showed lower somatic editing efficiency for target sites GW2T2 and GW7T6, while the efficiency of UPL3T11 editing remained the same (Figure 3b). The number of mutants identified in the M_1_ generation of plants inoculated with the pooled and individual BSMV-gRNAs showed similar trends (Figure 3c and d). The proportion of plants with mutations at the GW2T2, UPL3T11 and GW7T6 target sites dropped from 96% to 2.3%, from 27% to 19%, and from 51% to 21%, respectively. Only 5.7% of M_1_ plants carried mutations at two targets sites, and no mutants with all three targets edited were recovered (Figure 3c). These results suggest that with increase in the multiplexing level performed by simple pooling of BSMV-sgRNAs, we should expect decrease in the efficiency of editing of individual targets and the rate of multiplex gene editing.

**Figure 3.**
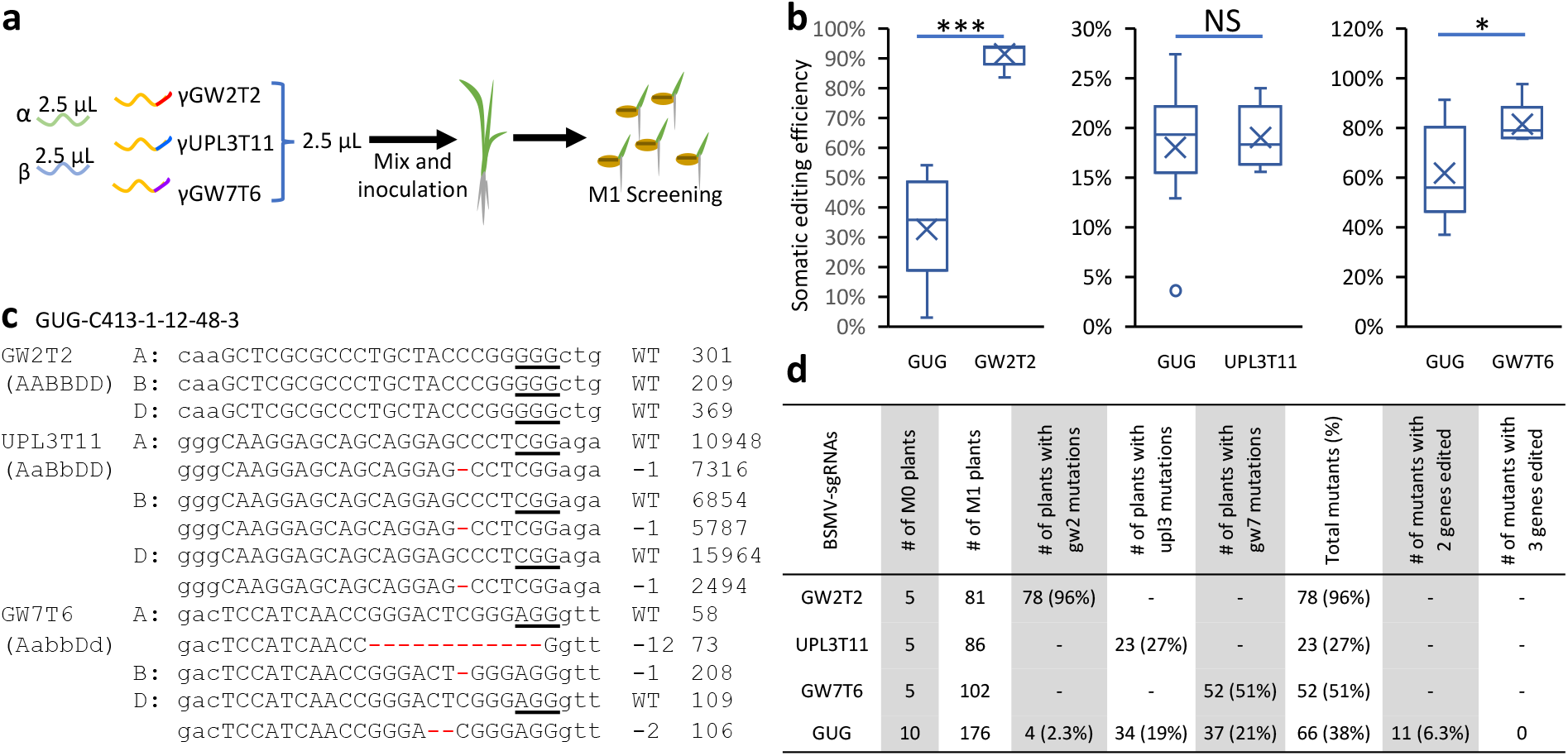
Multiplex gene editing by inoculating with the pooled BSMV-sgRNAs. **a)** A schematic pipeline of the BSMV-based multiplex editing using the sgRNA pool targeting *TaGW2* (GW2T2 sgRNA), *TaUPL3* (UPL3T11 sgRNA) and *TaGW7* (GW7T6 sgRNA). **b)** Comparison of somatic editing efficiency at each target site in plants inoculated with the BSMV-GUG pool or individual BSMV-sgRNAs. The data from all three genomes was combined to calculate the somatic editing efficiency. The multiplex and single target editing efficiencies were compared using the Student’s *t*-test; NS - *P* >0.05, * - *P* < 0.05, *** - *P* < 0.0001. **c)** Alignment of representative NGS reads from the M_1_ plants derived from the BSMV-GUG inoculated plants. The NGS reads were aligned to the A, B and D homoeologs of the *TaGW2, TaUPL3* and *TaGW7T6* genes. The deleted nucleotides are shown in red. The number of deleted nucleotides is shown on the right of the reads. The number of wild-type (WT) and mutated reads is shown on the right side of mutation types. The PAM sequences are underlined. **d)** The number of plants carrying mutations in one, two or all three genes obtained by inoculating with BSMV-GUG and the individual BSMV-sgRNAs.

Previously, the multiplexed genome editing strategy was used to generate variation in the promoters of genes controlling productivity traits (Rodríguez-Leal *et al*. 2017). Here, we tested the ability of the pooled BSMV-sgRNAs to edit the promoter of the *Q* gene (Figure 4a and Table S2), which controls a number of domestication traits in wheat linked with productivity (Simons *et al*. 2006). The Sanger sequencing of the 3 kb region upstream of the *Q* gene region revealed that cultivar Bobwhite has one SNP, one 160 bp deletion and one 160 bp insertion compared to the reference genome of cultivar Chinese Spring (The International Wheat Genome Sequencing Consortium (IWGSC) 2018) (Figure S3). The *Q* gene promoter was edited by inoculating the high-Cas9-expressing line C413 with a pool of five BSMV-sgRNA transcripts (henceforth, BSMV-pQT).

**Figure 4.**
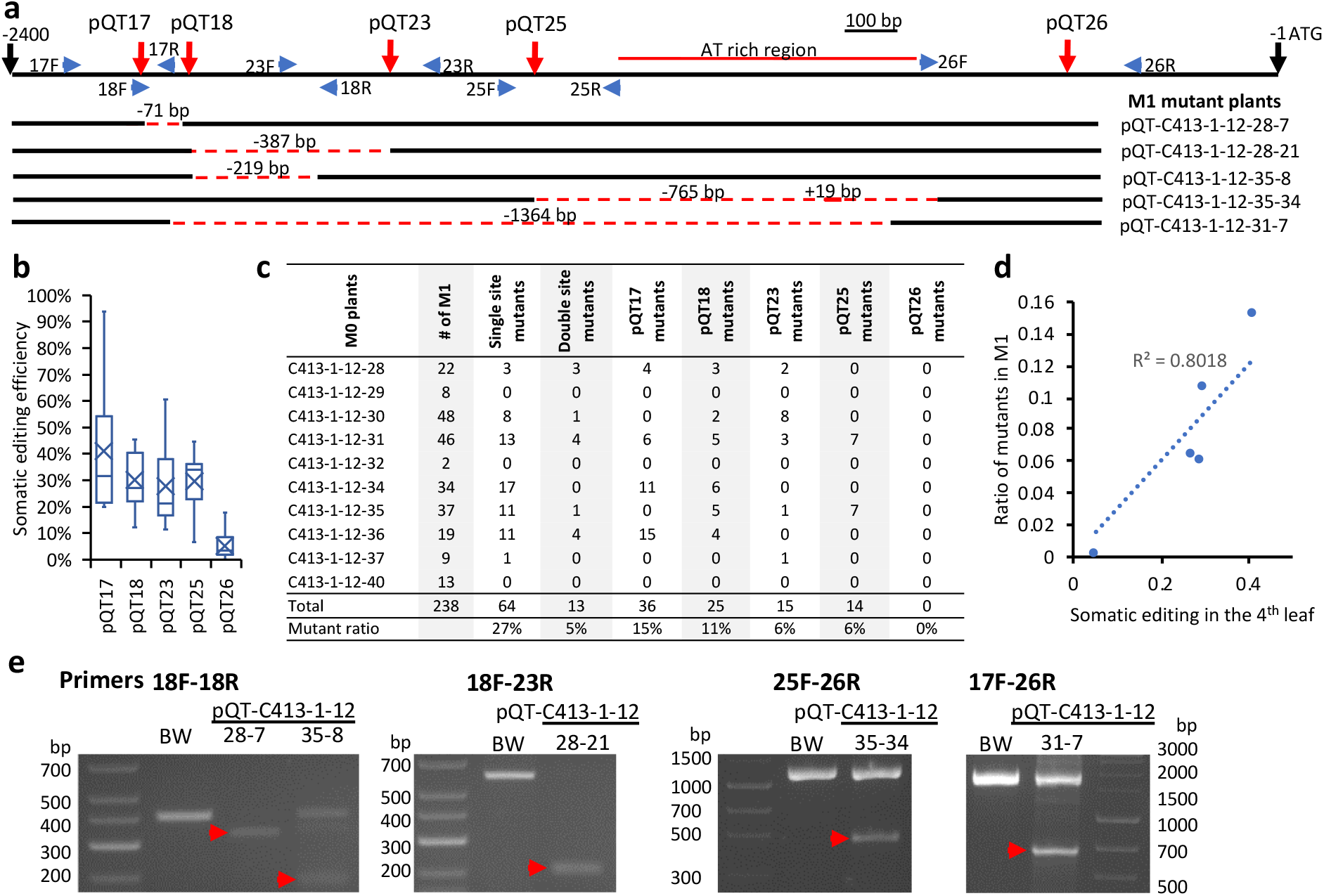
*Q* gene promoter editing using the pool of BSMV-sgRNAs (BSMV-pQT). **a)** The location of the BSMV-sgRNA target sites (red arrows) across the *Q* gene promoter region extending 2400 bp upstream of the start codon (ATG). The location of PCR primers used for genotyping are indicated by blue arrows. The location of an A/T rich region is shown by a red line. The deletions and insertions detected in five M_1_ plants are shown as red dashed and solid lines, respectively, with the sizes of deletions and insertions shown above. **b)** The somatic editing efficiency measured in the 4^th^ leaf (developed after inoculation) of plants inoculated with the BSMV-pQT pool. **c)** The summary of mutations discovered in the M_1_ generation plants inoculated with the BSMV-pQT pool. **d)** The relationship between the somatic editing efficiency and the proportion of mutants discovered in the M_1_ generation plants. Each data point stands for one BSMV-sgRNA target site. **e)** The results of PCR-based screening of the M_1_ generation plants for deletions in the promoter region of the *Q* gene promoter. A total of 11 pairs of primers (17F-18R, 17F-23R, 17F-25R, 17F-26R, 18F-18R, 18F-23R, 18F-25R, 18F-26R, 23F-25R, 23F-26R, and 25F-26R) were used in screening. The cultivar Bobwhite (BW) was used as wild-type control. The PCR fragments corresponding to deletions are shown by red arrows.

The means of somatic editing efficiency for each of the five target sites, including pQT17, pQT18, pQT23, pQT25, and pQT26, were 37%, 28%, 25%, 27% and 5%, respectively (Figure 4b). The proportions of M_1_ plants carrying heritable mutations at each target site were 15%, 11%, 6%, 6% and 0%, respectively (Figure 4c), and highly correlated with somatic editing efficiency (Figure 4d). About 32% of M_1_ plants carried heritable mutations in at least one target site. There were no M_1_ plants carrying heritable mutations at three or more targets. A total of 5% M_1_ plants had heritable mutations at two target sites, which is nearly 5 times lower than the proportion of M_1_ plants with mutations at a single target site (27%) (Figure 4c).

Further, we screened the edited plants for deletions spanning the regions between the pairs of the BSMV-sgRNA target sites. First, we tested M_0_ plants for deletions between target sites pQT17 and pQT18 by performing PCR using a pair of flanking primers 17F and 18R (Figure 4a). The NGS of amplicons showed that 5.2% - 19.9% of reads carried the 71 bp deletion expected upon successful targeting of both the pQT17 and pQT18 sites (Figure 4a, Table S3).

The *Q* gene promoter deletion screening in M_1_ plants was performed using 11 pairs of PCR primers. We identified five M_1_ plants that had deletions in the homozygous or heterozygous state or present as mosaic somatic mutations (Figure 4a and 4e). Two of these plants, pQT-C413-1-12-28-7 and pQT-C413-1-12-28-21, were homozygous for deletions between the pQT17 and pQT18 target sites and between the pQT18 and pQT23 target sites, respectively (Figure 4a). Compared to wild-type plants, these two deletion mutants did not exhibit visible differences in development or morphology. Consistent with this observation, the quantitative RT-PCR analysis of *Q* gene expression in the 3^rd^ and 6^th^ leaves at the 6-leaf developmental stage of M_1_ plants also did not detect significant changes compared to wild-type plants (Figure S4). Apparently, these deletions are located far away from the transcription start site and do not affect important regulatory regions controlling the *Q* gene expression and function. The plants with deletions in heterozygous state or present as non-fixed somatic mutations also not show visible phenotypic effects. Further studies aimed at creating additional deletions within the upstream regulatory region of the *Q* gene and more extensive phenotypic evaluation of created mutants are planned in the future.

### BSMV-sgRNA-based gene editing in the winter and spring wheat cultivars

Even though the BSMV-based method of sgRNA delivery method bypasses the wheat transformation and regeneration steps (Li *et al*. 2021), it still depends on the availability of a wheat line expressing the Cas9 gene. In spite of recent advances in the wheat regeneration methodology (Debernardi *et al*. 2020), the development of transgenic wheat cultivars remains a time-consuming process and has varying levels of success. To broaden the range of wheat cultivars amenable to editing using the BSMV-based sgRNA delivery system, we transferred the highly expressed allele of Cas9 from line 707 into elite spring wheat line 3613474 from CIMMYT and winter wheat line KS080093K-18 from Kansas wheat breeding program (Figure 5a and 5b). The backcrossed progeny of both lines, now carrying the highly expressed allele of Cas9, was inoculated with the RNA transcripts synthesized from the BSMV-sgRNA constructs targeting the *Q* and *TaGW7* genes. Similar to the results obtained by inoculating the high-Cas9 C413 line with BSMV-QT1 (Figure 1), the BC_2_ F_3_ plants from the 3613474 family inoculated with BSMV-QT1 also exhibited spear-shaped speltoid spikes suggestive of the presence of the loss-of-function mutations in the *Q* gene. Consistent with this result, the high level of somatic editing (97%) was observed for the QT1 target in these plants (Figure 5c). In the M_1_ generation, 61% of plants carried mutations in at least one homoeologous copy of the *Q* gene with nearly 30% of plants carrying mutations in all three homoeologs of the *Q* gene (Figure 5d). The plants from the 3613474 and KS080093K-18 backcross families inoculated with BSMV-GW7T6 showed 59% and 64% somatic editing efficiency, respectively (Figure 5c). In the M_1_ generation, 11% and 17% of 3613474- and KS080093K-18-derived plants inherited mutations in the *TaGW7* gene, respectively (Figure 5d).

**Figure 5.**
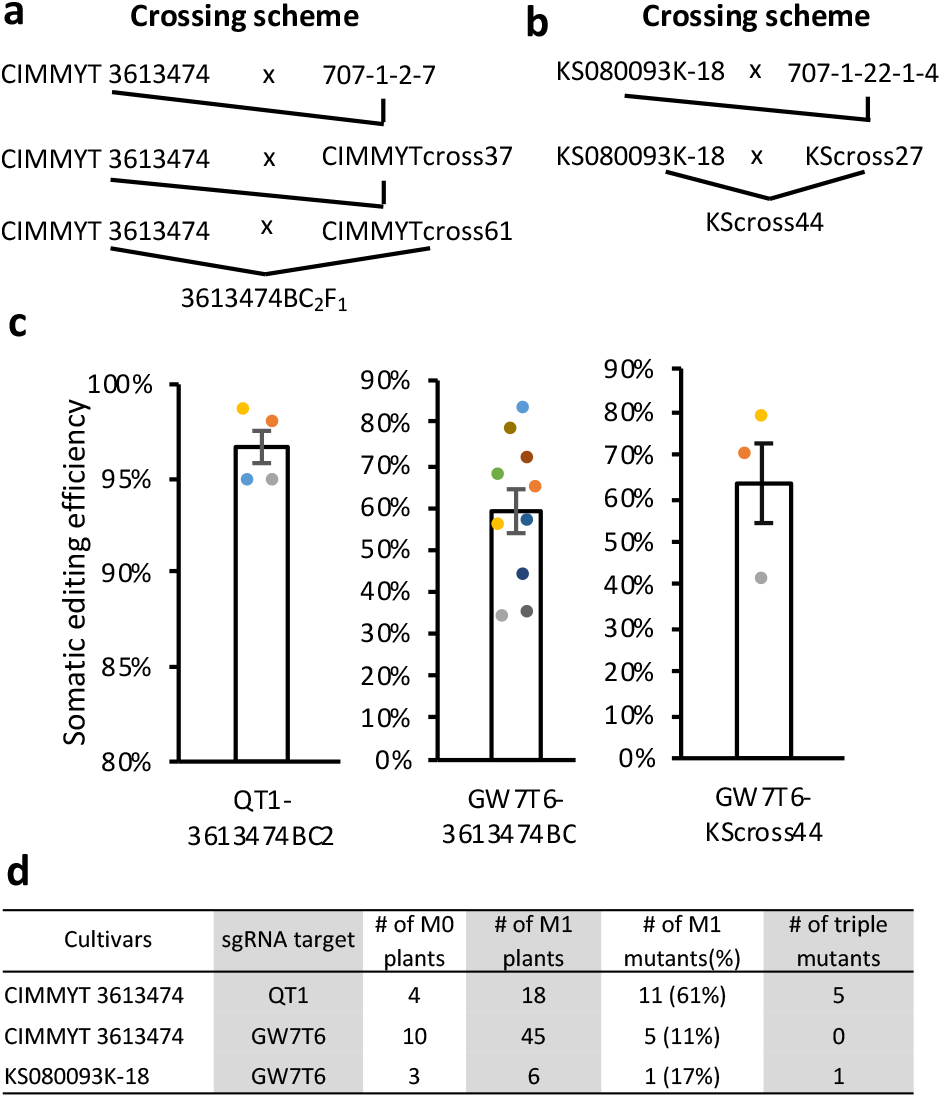
BSMV-sgRNA-based gene editing of the spring and winter cultivars. Introgression of the highly expressed Cas9 allele from the high-Cas9 707 line (Figure S1) into **a)** CIMMYT line 3613474 (spring wheat) and **b)** Kansas line KS080093K-18 (winter wheat). The BC2F3 and BC1F2 lines derived from the crosses with 3613474 and KS080093K-18, respectively, were used for BSMV inoculation. **c)** The somatic editing efficiency was estimated 1) for BSMV-QT1 and BSMV-GW7T6 constructs used to inoculate lines derived from 3613474 and 2) for BSMV-GW7T6 used to inoculate lines derived from KS080093K-18. The editing efficiency was evaluated in the 4^th^ leaf of inoculated plants. The data from all three genomes are combined to calculate the editing efficiency. **d)** The summary of the M_1_ generation mutants obtained by the BSMV-sgRNA-based editing of wheat cultivars expressing the introgressed high-Cas9 allele.

## Discussion

Our study shows that the efficiency of the BSMV-based editing and the chance of obtaining heritable mutations depend on the sgRNA design and the levels of Cas9 expression. This dependency holds for both single BSMV-sgRNA and multiplexed BSMV-sgRNA constructs, indicating that the development of the transgenic wheat line with the adequate levels of Cas9 expression is critical for successful BSMV-sgRNA-based editing. Once a high-Cas9 line is developed, it can be used either directly in gene editing experiments or as a donor of the high-expressing Cas9 allele for introgression into other cultivars.

We showed that somatic editing frequency in the M_0_ plants inoculated with the BSMV-sgRNA transcripts correlates with the frequency of heritable mutations in the M_1_ generation. It appears that in the BSMV-sgRNA-based gene editing experiments we can skip the step of protoplast-based testing of the sgRNA editing efficiency (Wang *et al*. 2019), and instead directly assess the sgRNA editing efficiency by sequencing the DNA extracted from the newly grown seedling leaves of the BSMV-sgRNA inoculated plants. Such early assessment of somatic editing efficiency can help to quickly adjust on-going experiments. For example, in case of discovering that a critical sgRNA has low editing efficiency, the number of the BSMV-sgRNA inoculated plants could be increased to increase heritable mutation recovery rate or sgRNA could be re-designed to improve its efficiency in the follow up experiments.

A previous study (Li *et al*. 2021) show that the multiplex wheat genome editing is possible with the pooled BSMV-sgRNAs. Our results indicated that the level of multiplexing for achieving simultaneous editing of all targets could be limited. The likelihood of recovering mutations at multiple sites across a genome appears to be the product of editing efficiency of individual targets. Probably this happens because the chance for the same plant cell to be infected by multiple BSMV particles with distinct guides is reduced with increase in the level of multiplexing. For effective editing of multiple targets using the BSMV-sgRNA-based method, we should explore alternative strategies. One of the possible approaches is to create BSMV-sgRNA constructs expressing an array of multiple guide RNAs simultaneously. However, the level of multiplexing in this case will likely be limited by the restrictions on the size of a genome that could be packed into a viral particle. The usage of both the β- and γ-chains of the BSMV genome for subcloning the sgRNAs or development of the BSMV-sgRNA system based on the Cas12a editors (Zetsche *et al*. 2015; Wang *et al*. 2021), which utilize much shorter guides, are two paths to explore for expanding the multiplexing capacity of the BSMV-sgRNA-based editing system.

The utility of CRISPR-induced *cis*-regulatory diversity for improving productivity traits was clearly demonstrated in tomato (Rodríguez-Leal *et al*. 2017). In our study, we showed that the multiplexed BSMV-sgRNA-based system could be an effective tool for creating deletions in the upstream regulatory region of the *Q* gene. The approach was highly effective and allowed us to quickly recover deletion variants in the next generation of the BSMV-sgRNA-inoculated wheat lines. The distribution of editable sites that could be targeted by Cas9 in the *Q* gene promoter did not allow us to create deletions near the transcription start site and recovered mutants did not show significant changes in the gene expression level or phenotype. Nevertheless, our results demonstrate that this editing strategy opens opportunities for exploring novel *cis*-regulatory diversity in the wheat genome and modulating regulatory effects of transcription factors controlling agronomic traits. One of the shortcomings of pooling multiple BSMV-sgRNAs was the reduction in the efficiency of individual target editing, which resulted in relatively low frequency of any given deletion variant. To increase the chance of recovering specific deletions using the BSMV-sgRNA-based system, the reduced levels of multiplexing should be considered (e.g., pooling only two BSMV-sgRNAs targeting sites flanking the region of interest).

We showed that we could perform editing of other wheat cultivars by introgressing the high expressing allele of Cas9 using marker-assisted forward selection. The lines developed in our study could serve as donors of the high-Cas9 alleles capable of maintaining the effective BSMV-sgRNA-induced genome editing. This strategy could be applied for the functional analyses of candidate genes or generation of desired mutations in any genetic backgrounds and essentially removes the cultivar-specific constraints on targeted genetic modification of the wheat genome. Contrary to approaches that are based on random genome-wide chemical mutagenesis of thousands of lines (Henry *et al*. 2014; Krasileva *et al*. 2017), these functional screens can be used to analyze the effects of mutations without the confounding impact of mutations in other parts of the genome. As soon as the donors of high-expressing Cas9 lines and a virus-based CRISPR delivery system are available (Yin *et al*. 2015; Ellison *et al*. 2020; Li *et al*. 2021), a similar gene editing approach could be used for modifying the genomes of other crops. Considering the ease with which the BSMV-sgRNA constructs can be designed to target hundreds of targets and the simplicity of BSMV-sgRNA inoculation procedure this approach provides an opportunity for establishing the high-throughput functional screens in genetic studies to identify causal mutations or novel variants beneficially affecting agronomic traits.

## Methods

### Plasmid construction

The previously reported plasmids (Scofield *et al*. 2005) with the α, β and γ chains of the ND18 strain of barley stripe mosaic virus (BSMV), referred to as pBSMVα, pBSMVβ and pBSMVγ, were used in our study (Figure 1a). The virus induced gene silencing (VIGS) construct pBSMVγPDS previously developed for targeting the barley phytoene desaturase gene (*HvPDS*) was used in our study to silence *TaPDS* in wheat (Holzberg *et al*. 2002; Scofield *et al*. 2005) (Figure 1a). For delivering sgRNAs into plant cells using BSMV, the sgRNA sequences were inserted into the pBSMVγPDS plasmid after removing the *PDS* fragment by PacI and NotI digestion (New England BioLabs, Ipswich, MA, catalog numbers R0547 and R0189). The sequences targeted in our study are listed in Table S2. All the primers, DNA oligos and double stranded DNA fragments are synthesized by Integrated DNA Technologies (Coralville, IA, USA) (Table S4). PCR reactions were performed using NEBNext® High-Fidelity 2X PCR Master Mix (New England BioLabs, catalog number M0541) following the manufacture’s protocol unless specified differently. For example, to insert the sgRNA for target QT1 into pBSMVγPDS, the sgRNA was amplified from pU6sg-QT1 construct (Wang *et al*. 2016) using a primer pair targetVIGSsF and targetVIGSsR. The PCR fragments and the plasmid pBSMVγPDS were digested using PacI and NotI, the PCR fragment and the plasmid fragment without *PDS* gene were isolated from agarose gel using QIAquick Gel Extraction Kit (Qiagen, catalog number 28706). The digested PCR products and the pBSMVγ plasmid were ligated using T4 ligase (New England BioLabs, catalog number M0202). This plasmid was named pBSMVγQT1 (Figure 1b). All inserted DNA fragments were validated by Sanger sequencing. All the Sanger sequencing in this study were done using BigDye™ Terminator v3.1 Cycle Sequencing Kit (ThermoFisher Scientific, catalog number 4337455).

To test the effect of mobile RNAs on BSMV-sgRNA-induced gene editing, the fragments including GW2T2-sgRNA and mobile RNAs were synthesized at Integrated DNA Technologies (Table S4). The mobile RNAs included the coding sequences of *FT* gene from *Arabidopsis* (Ellison *et al*. 2020) and its wheat ortholog *Vrn3* (Yan *et al*. 2006), and wheat methionine and isoleucine tRNAs tRNA^Met^ and tRNA^Ile^ (Figure 2a). The GW2T2-sgRNA without mobile elements were amplified from the previously developed pBUN421-GW2T2 plasmid (Wang *et al*. 2018a) using the primer pair VIGS-GW2T2sgRNA-F and VIGS-sgRNA-R. Both the synthesized DNA fragment and the PCR products were subcloned into PacI- and NotI-digested pBSMVγPDS using NEBuilder^®^ HiFi DNA Assembly Master Mix (New England BioLabs, catalog number E2621) following the manufacture’s protocol.

The UPL3T11-sgRNA with flanking sequences overlapping with the PacI and NotI digested pBSMVγPDS (Table S4) were synthesized at Integrated DNA Technologies. The sgRNAs with flanking sequences for GW7T6, pQT17, pQT18, pQT23, pQT25 and pQT26 were obtained by amplifying the sgRNA scaffold of the pBUN421 plasmid (Xing *et al*. 2014) using the forward primers paired with reverse primer VIGS-sgRNA-R (Table S4). Both synthesized DNA fragments and PCR products were subcloned into PacI- and NotI-digested pBSMVγPDS using NEBuilder^®^ HiFi DNA Assembly Master Mix.

### Sequencing of the *Q* gene promoter in cv. Bobwhite

Three pairs of primers (Table S4) targeting overlapping genomic regions were used to amplify the *Q* gene promoter in cv. Bobwhite. The PCR fragments of primer pair pQT4MiF - pQT17MiR and primer pair pQT17MiF - pQT25MiR were subcloned into pCRblunt using Zero Blunt™ TOPO™ PCR Cloning Kit (ThermoFisher Scientific, catalog number 450245) following the manufacture’s protocol. For each subcloned fragment, three colonies were sequenced. The PCR product of primer pair pQT25checkF - Q5endR3 was sequenced directly without subcloning.

### Plants and growth conditions

A transgenic line 7438 expressing previously reported wheat-codon-optimized Cas9 in the background of cv. Bobwhite (Wang *et al*. 2016) was utilized for the initial testing of BSMV-sgRNA-based gene editing. Later, to improve the BSMV-based gene editing efficiency, additional transgenic lines in the background of cv. Bobwhite were screened to identify plants with the high levels of Cas9 expression (high-Cas9). The 707 and C413 lines expressing maize-codon-optimized Cas9 at high level were used for further research (Figure S1). The 707 line was used as a donor of high-Cas9 allele for intogression into CIMMYT spring wheat 3613474 and Kansas winter wheat KS080093K-18. These two lines were crossed with 707, and then Cas9 positive F_1_ plants were selected by PCR using the primer pair zCas9-F and zCas9seq1 (Table S4) and backcrossed one and two times to KS080093K-18 and 3613474, respectively. The Cas9-positive lines from the backcrossed plants were used for BSMV-sgRNA inoculation.

All plants were grown in ¼ litter square pots filled with the Berger BM1 growing mix (Hummert, USA, catalog number 10120000) in a growth chamber at 24 °C day/20 °C night conditions with 16 hour supplemental light. The plants were grown in ¼ litter square pots for two weeks after inoculation, and then transplanted into 1 L square pots. The M_1_ generation plants were grown in the 128-well plastic trays filled with the Berger BM1 nutrient retention soil.

### In vitro transcription of BSMV-sgRNA constructs and plant inoculation

The α, β and γ virus chains were transcribed *in vitro* from plasmids pBSMVα, pBSMVβ and pBSMVγ linearized by digesting with MluI (New England BioLabs, catalog number R3198S), SpeI (New England BioLabs, catalog number R0133S) and MluI enzymes, respectively. The pBSMVγPDS plasmid and all plasmids expressing BSMV γ chain carrying sgRNAs were linearized with BssHII (New England BioLabs, catalog number R0199S). In vitro transcription was performed in 20 µL reaction volume using HiScribe™ T7 High Yield RNA Synthesis Kit (New England BioLabs, catalog number E2040S) following the Capped RNA Synthesis protocol provided by the manufacturer. The m7G(5’)ppp(5’)G RNA Cap (New England BioLabs, catalog number S1404L) was used as Cap Analog with 4:1 of Cap Analog:GTP ratio. The quality and concentration of RNA transcripts (usually within 2-2.5 µg/µL range) were assessed on agarose gel.

The second leaf of wheat seedlings at the two-leaf stage was inoculated with the FES buffer, BSMV, BSMV-PDS or BSMV-sgRNAs. Each plant was inoculated using the mixture of 60 µL FES buffer and transcription products of BSMV α, β and γ chain (2.5 µL each). The FES buffer includes 7.51 g/L glycine, 10.45 g/L K_2_ HPO_4_ dibasic, 10 g/L sodium pyrophosphate decahydrate, 10 g/L bentonite and 10 g/L celite. The inoculation was performed by hand-rubbing the 2^nd^ leaf from the base to the tip while wearing clean nitrile gloves. The procedure was repeated 3 times for each plant, each time applying 20 µL of the mixture. Immediately after inoculation, the plants were covered by plastic bags to keep the moisture. Plastic bags were removed 5 to 7 days after inoculation. In each experiment, three groups of controls including FES buffer, wild-type BSMV and BSMV-PDS were used. The virus infection symptoms were usually observed 7 days after inoculation, while the PDS RNAi phenotype appeared 12 days after the inoculation.

### Genotyping of BSMV inoculated plants and screening of mutants in M1 generation

To detect target editing events, a 2 cm-long segment of the 4^th^ leaf was sampled about two weeks after inoculation when BSMV-PDS-inoculated control plants showed photobleached leaf symptoms. DNA was isolated using the TPS buffer (100 mM Tris-HCl, 10 mM EDTA, 1 M KCl, pH8.0) as described (Wang *et al*. 2021). The gene editing efficiency in the M_0_ and M_1_ plants was assessed using the next generation sequencing (NGS) approach, as previously described (Wang *et al*. 2018a).

The deletions in the *Q* gene promoter region were detected using various combinations of PCR primers (Figures 4a and 4e, Table S4) listed in. The examples of agarose gel images with PCR products showing evidence of deletions in the *Q* gene promoter are shown in Figure 4.

### RNA isolation and qPCR

RNA isolation was performed using the TRIZOL reagent (ThermoFisher Scientific, catalog number 15596026) following the manufacture’s protocol. The cDNA was obtained by reverse transcription using SuperScript™ III First-Strand Synthesis SuperMix for qRT-PCR (ThermoFisher Scientific, catalog number 11752050). The Cas9 expression in the transgenic plants was measured by qRT-PCR using the primer pair zCas9-F and zCas9seq1 and RNA isolated from the 2^nd^ leaf of two-week-old seedlings. The *Q* gene expression in the M_1_ plants was measured by isolating RNA from the 3^rd^ and 6^th^ leaf at 6-leaf development stage. The samples were frozen in liquid nitrogen immediately and then stored under – 80 °C until RNA was isolated. The qPCR reaction was performed using the PowerUP SYBR Green Master Mix (ThermoFisher Scientific, catalog number A25741) following the manufacture’s protocol. NEBNext® High-Fidelity 2X PCR Master Mix (New England BioLabs, catalog number M0541) and SybrGreen were used to assess the expression level of the Q gene on chromosome 5A. The specificity of the Q gene primers, Q5ArtF4 and Q5ArtR4 (Table S4), was validated using nullisomic-tetrasomic genetic stocks (Figure S5). The *TaActin* gene expression was used as reference.

## Author Contributions

W.W. designed and conducted gene editing experiments, genotyped and phenotyped transgenic plants, collected and analyzed phenotyping data, and helped with drafting the manuscript; Z.Y. conducted gene editing experiments; F.H. contributed to the analysis of NGS data; G.B. performed Sanger sequencing; H.T. coordinated biolistic transformation part of the project; A.A. designed and conducted NGS experiments; E.A. conceived idea, interpreted results, coordinated project, and wrote the manuscript. All authors read the manuscript and approved the final version.

## Acknowledgements

This project was supported by the Agriculture and Food Research Initiative Competitive Grants 2021-67013-34174 and 2020-67013-30906 from the USDA National Institute of Food and Agriculture, and grants from the Bill and Melinda Gates Foundation (INV-004430) and Kansas Wheat Commission. We thank Alyssa Dunnivan for help with DNA extraction and preparation of PCR amplicons for NGS analysis and Dwight Davidson for the greenhouse management and phenotyping data collection.

## Supplementary Information

**Figure S1**. The expression level of Cas9 in transgenic plants. The quantitative PCR of Cas9 was conducted using Cas9F and Cas9seq1 (Table SM1). The *TaActin* gene was adopted as reference. The relative expression level of Cas9 is shown as means ± SE based. The T3 progeny of line 7438 and line C413 were used, the T4 progeny of line 707 were used. The biological replicate number for line 7438, C413 and 707 are 6, 9 and 4, respectively. Three biological replicates were used for cv. Bobwhite in both bar plots.

**Figure S2**. Relationship between the mutagenesis ratio in the M1 progeny of plants inoculated by BSMV-GW2T2 (a, b, and d) or BSMV-GW2T2 (c and e) and the somatic editing efficiency evaluated in the 6^th^ (a), 4^th^ (b and c), and 2^nd^ (d and e) leaf. The 2^nd^ and 4^th^ leaf were sampled at 4-leaf stage. The 6^th^ leaf was sampled at 6-leaf stage. Each data point represents an individual plant.

**Figure S3**. Alignment of Q gene promoter from cultivar Chinese Spring and Bobwhite. The top sequence is Chinese Spring, and the bottom sequence is cultivar Bobwhite. The promoter region is 2990 bp long upstream of the start codon of Q gene in both cultivars. The missed nucleotides in the alignment are shown as “-”; the mismatch in the alignment is shown in red fonts. The position of the last nucleotide in each row is shown on the right side of the alignment.

**Figure S4**. Expression of *Q* gene A genome allele in the M1 plants at 6-leaf stage. The A genome specific primers, Q5ArtF4 and Q5ArtR4 (Table S4), were applied. *TaActin* gene was used as reference. The 3^rd^ and 6^th^ leaf in M1 plants at 6-leaf stage were sampled for RNA isolation followed by reverse transcription to get cDNA. The plant pQT-C413-1-12-28-7 and pQT-C413-1-12-28-21 carrying homozygous long deletions are highlighted with red box in the bar plots. All the rest of the plants have wild type alleles all the promoter of Q gene. The results are shown as mean ± SE based on three technical repeats.

**Figure S5**. Validation of the *Q* gene A genome specific primers for RT-PCR. PCR amplification of cv. Bobwhite, cv. Chinese Spring (CS) and nullisomic-tetrasomic lines using the genome-specific primer Q5ArtF4 and Q5ArtR4 (Table S4) was performed. DNA isolated from six nullisomic-tetrasomic lines (N5A-T5D, N5B-T5A, N5DT5A, and N5DT5B), cv. Chinese Spring (CS), and cv. Bobwhite (BW). NTC is no template control of PCRs.

**Table S1**. The efficiency of editing based on the BSMV-sgRNA delivery system in transgenic wheat lines with the low (7438) and high (C413) levels of Cas9 expression.

**Table S2**. The target sites selected for BSMV-sgRNA-based editing in this study.

**Table S3**. Proportion of NGS reads carrying 71-bp deletion.

**Table S4**. The primers, oligos and synthesized double-strand DNA used in this study.

