## Supplementary Information for "Multiplexed promoter and gene editing in wheat using the virus-based guide RNA delivery system"

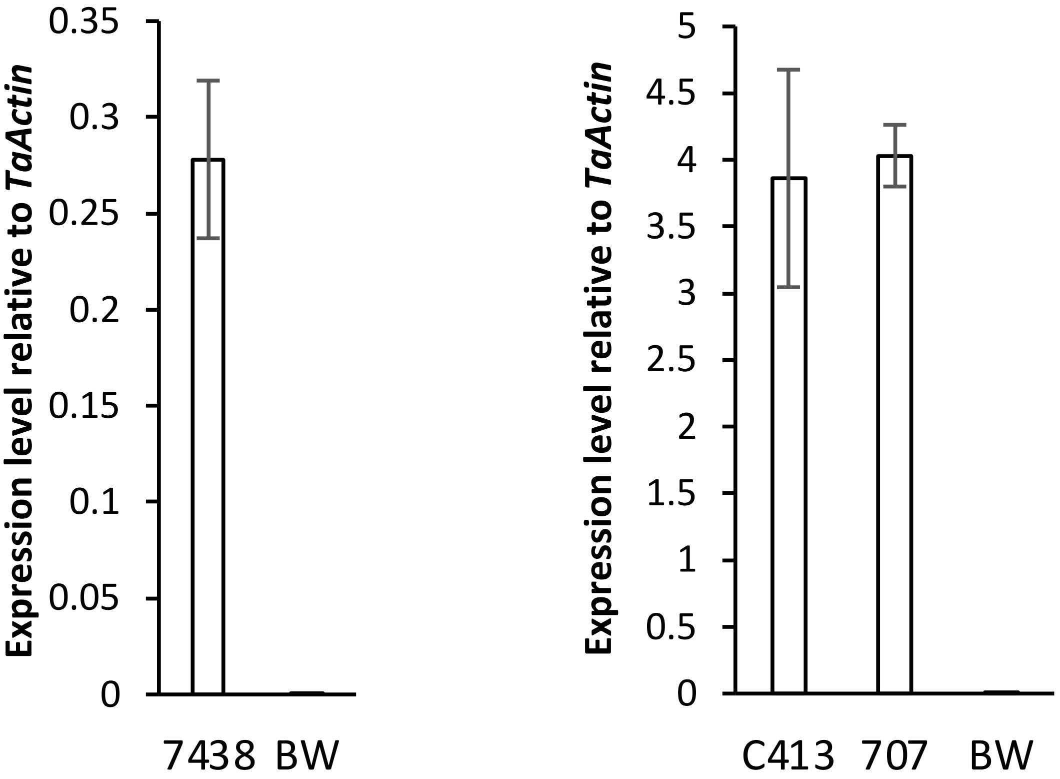


**Figure S1.** The expression level of Cas9 in transgenic plants. The quantitative PCR of Cas9 was conducted using Cas9F and Cas9seq1 (Table SM1). The *TaActin* gene was adopted as reference. The relative expression level of Cas9 is shown as means ± SE based. The T3 progeny of line 7438 and line C413 were used, the T4 progeny of line 707 were used. The biological replicate number for line 7438, C413 and 707 are 6, 9 and 4, respectively. Three biological replicates were used for cv. Bobwhite in both bar plots.

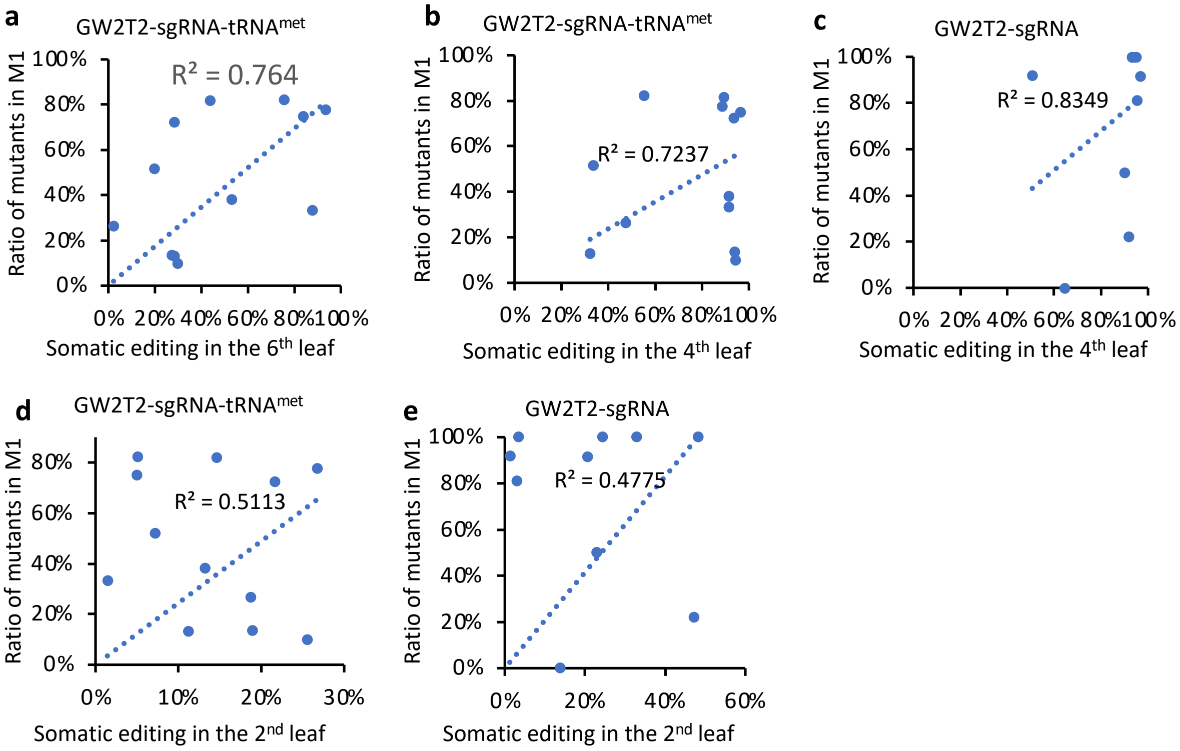


**Figure S2.** Relationship between the mutagenesis ratio in the M1 progeny of plants inoculated by BSMV-GW2T2 (a, b, and d) or BSMV-GW2T2 (c and e) and the somatic editing efficiency evaluated in the 6th (a), 4th (b and c), and 2nd (d and e) leaf. The 2nd and 4th leaf were sampled at 4-leaf stage. The 6th leaf was sampled at 6-leaf stage. Each data point represents an individual plant.

```
TAAATCATGGTAGTTACATCTCGTCATATTCTATGAACAATCATATTTCTTATGAAATCTTGCACAACTATCACGCCATAGCATGACCGACACATAACTA 100
TAAATCATGGTAGTTACATCTCGTCATATTCTATGAACAATCATATTTCTTATGAAATCTTGCACAACTATCACGCCATAGCATGACCGACACATAACTA 100

TAAGACACTATTTTTCTTTCAAAAGCTTGATAGTACTTTCTTCAAAAGCTCATACTACTACTAAACACGAGCAAGTCTTGACCAAGATGAAGCGCGCGCG 200
TAAGACACTATTTTTCTTTCAAAAGCTTGATAGTACTTTCTTCAAAAGCTCATACTACTACTAAACACGAGCAAGTCTTGACCAAGATGAAGCGCGCGCG 200

GGTGTAGCAAGGCTCCGTTTTCGTAGTAAAAACCATCTCCTTTTCTCGTTCCATCTTTTCTCAACCCGTCAAATCAAGCGAAAGCAATACTAGAACAGCG 300
GGTGTAGCAAGGCTCCGTTTTCGTAGTAAAAACCATCTCCTTTTCTCGTTCCATCTTTTCTCAACCCGTCAAATCAAGCGAAAGCAATACTAGAACAG-- 298

CTGCTATACACACGATGGTGTGTGGACGATTCCTGAACGATAACGCAGATCAACTCATCCATGCACAACAGTAAAGTGAAGCAGACGGCTGGATTTGGCC 400
---------------------------------------------------------------------------------------------------- 298

GTCGTCTCCTTGTCGTGCACGCAGTATCGTCTTTACAGCAGTTTCGATACTAGAACAGAACCATGTGCTCCAGTACGCAACAAGCGAAGCCATCGCGACA 500
----------------------------------------------------------AACCATGTGCTCCAGTACGCAACAAGCGAAGCCATCGCGACA 340

AAAACAAAATGAAGAAAAAGAACGGTGCGCTCCACCTCCCCCTCCGAGGCTCCGACCCGCCCGCCCGCAGCTCAGCTCAAGGCGCGCACACAGACACAGG 600
AAAACAAAATGAAGAAAAAGAACGGTGCGCTCCACCTCCCCCTCCGAGGCTCCGACCCGCCCGCCCGCAGCTCAGCTCAAGGCGCGCACACAGACACAGG 440

CCGGAGGGGGGCGTTCGGGCCGGCACGTACCCCCGCCCCACGTCCCGATCACCGGGTCGCCTCACCCTCACGGGCTCGCTCTCAACCGGTCAACGTACTGT 700
CCGGAGGGGGGCGTTCGGGCCGGCACGTACCCCCGCCCCACGTCCCGATCACCGGGTCGCCTCACCCTCACGGGCTCGCTCTCAACCGGTCAACGTACTGT 540

ACCGCACCGGTGCAGCCCATTTACGGCGCTGTCGCGGTGTCGCCGTCCTCCCTCCGTCCGTCCATTCCATCGGGTCCTCCCGTGGCCGTGCCGAGCCCCC 800
ACCGCACCGGTGCAGCCCATTTACGGCGCTGTCGCGGTGTCGCCGTCCTCCCTCCGTCCGTCCATTCCATCGGGTCCTCCCGTGGCCGTGCCGAGCCCCC 640

ACGCGGTGCCGTGACCGCGACCGTACGCGAGCCCGCCCGCCCGGCCGGCCCGTGACCACGAGCGTAAGGTTACGAGGATGCTACATACATGCGCGTACGG 900
ACGCGGTGCCGTGACCGCGACCGTACGCGAGCCCGCCCGCCCGGCCGGCCCGTGACCACGAGCGTAAGGTTACGAGGATGCTACATACATGCGCGTACGG 740

CTTACGTAGACAGAGCGAGTATACATGGCAGACTTTGGCGCGGGTGGGCGTCTCTCGGGTGGTGGGGGATGCATTTGTCGAAAGGAACAACGTTTCCGAT 1000
CTTACGTAGACAGAGCGAGTATACATGGCAGACTTTGGCGCGGGTGGGCGTCTCTCGGGTGGTGGGGGATGCATTTGTCGAAAGGAACAACGTTTCCGAT 840

GGGGCGAGCGGGGGCGTGACCGGTCGACCGGAACGGCACAGTGGCCCGGACAGGTACGCCTGTGGCTGCGTACCCGCCCTCGCGTGGGTCTACGTCGAAT 1100
GGGGCGAGCGGGGGCGTGACCGGTCGACCGGAACGGCACAGTGGCCCGGACAGGTACGCCTGTGGCTGCGTACCCGCCCTCGCGTGGGTCTACGTCGAAT 940

AATTGCAC-------------------------------------------------------------------------------------------- 1108
AATTGCACGAGCGCTGCTATGCACACGACGCATAGCAGACGATTTGTGCACGATAGGGCATATCCGTGCGTCCGTGCATATTATCAAAGTGCAGCAGACG 1040

--------------------------------------------------------------------CCATCCCATTACACCCGGGCCCCGGCGAGCAA 1140
GCTAGATGAAAATCGTGCACCTGTCGTCCATGGAGTGCATCGTCTGCATAGCTGTTTCGTAATTGCACCCATCCCATTACACCCGGGCCCCGGCGAGCAA 1140

AACAGTACCCGGACCTAGCCTGCAACCCCCAAGCCCCCGACGCGGTACATGCACGCATACACACACACGAAAGAGAGCGGTGCACGCAGTACGTACACAC 1240
AACAGTACCCGGACCTAGCCTGCAACCCCCAAGCCCCCGACGCGGTACATGCACGCATACACACACACGAAAGAGAGCGGTGCACGCAGTACGTACACAC 1240

CGGCAGGCGGACGCGGTACGTGCTAGGCTAGGCCAGGCTAGATTGGTCCAGCTGCCGGCTCCCCCCGTGCTCTCGTCGGCACGGCGCGACATGCCGTCAC 1340
CGGCAGGCGGACGCGGTACGTGCTAGGCTAGGCCAGGCTAGATTGGTCCAGCTGCCGGCTCCCCCCGTGCTCTCGTCGGCACGGCGCGACATGCCGTCAC 1340

GTACCTGCTCCGCCTGTGGCCCTTGGCGCTTGCGCCCGGCCGGCCGCGGTCGGGTCAACACACGAGGCCTCCCAGATCGGGCGCGGCATGCATGTGCCGC 1440
GTACCTGCTCCGCCTGTGGCCCTTGGCGCTTGCGCCCGGCCGGCCGCGGTCGGGTCAACACACGAGGCCTCCCAGATCGGGCGCGGCATGCATGTGCCGC 1440

CGGTACGTATGTACGTATACCGGCGCGGATTAATTTAGAGTTCGATTTGATTAGAGGGAGGGAGGGGGGCGTGCGGTGCACTTGGGCAATGTAATGCGGT 1540
CGGTACGTATGTACGTATACCGGCGCGGATTAATTTAGAGTTCGATTTGATTAGAGGGAGGGAGGGGGGCGTGCGGTGCACTTGGGCAATGTAATGCGGT 1540

CCTGCGAGGAGGGATCTCATCTAACCTAGCAGCACAGGGCGTACGGCCGGGGGCTTATCTTACTCTCGCTAGGTGCCTAAGATAACCAGCATGAGTTGAT 1640
CCTGCGAGGAGGGATCTCATCTAACCTAGCAGCACAGGGCGTACGGCCGGGGGCTTATCTTACTCTCGCTAGGTGCCTAAGATAACCAGCATGAGTTGAT 1640

GGTGCCGGCCGTTAACAATTCCAACCAAAAGCTAATCGTCTGGTGCACACGCAATGGTGGACACTATCATACCATGGATCACGTGGGTGGTTCTTTGTCC 1740
GGTGCCGGCCGTTAACAATTCCAACCAAAAGCTAATCGTCTGGTGCACACGCAATGGTGGACACTATCATACCATGGATCACGTGGGTGGTTCTTTGTTC 1740

ATGCCACTGGTTTTTCCTCTTTCGCTAGAGTTAATTAAGCGAATGGCTTTTGAATCCGTGTTTGTTCATGTCGGTCAAATCAAATCTCATGAATATGTTG 1840
ATGCCACTGGTTTTTCCTCTTTCGCTAGAGTTAATTAAGCGAATGGCTTTTGAATCCGTGTTTGTTCATGTCGGTCAAATCAAATCTCATGAATATGTTG 1840

TACCGTTTGTCCTAGATGGAGCTGGATTAAATAATTTCGGCATTTGTGGGTGCCCCTCCATACGATGTACTGGAGAGGGTTTTAAAACATTTTGTTGCTT 1940
TACCGTTTGTCCTAGATGGAGCTGGATTAAATAATTTCGGCATTTGTGGGTGCCCCTCCATACGATGTACTGGAGAGGGTTTTAAAACATTTTGTTGCTT 1940

AGCGTTTGGCTAGCGATGTAATAACAAAATAGTGTCATGACATCTGCCACAAAGGTGCGTCGACATCCATACTTTACTCTTCGTTGCACTACAAACATAT 2040
AGCGTTTGGCTAGCGATGTAATAACAAAATAGTGTCATGACATCTGCCACAAAGGTGCGTCGACATCCATACTTTACTCTTCGTTGCACTACAAACATAT 2040

TTTGTGATGTACGCTCCGTGTGATATATATACTTGTATCTTTAGAAAGGCCTAGAGTCTTAGGTGTAAGACCTTTCTAGCTGCAGGCCTGATTCGCACAA 2140
TTTGTGATGTACGCTCCGTGTGATATATATACTTGTATCTTTAGAAAGGCCTAGAGTCTTAGGTGTAAGACCTTTCTAGCTGCAGGCCTGATTCGCACAA 2140

GGTGCTACTTGGTTGTTGCATATAAAAAACGGTAATACTATAATACTCTGTCCCTCCGGGACTAAATTACTTAGCAAGTTGACATTTGTTTATATTTAGC 2240
GGTGCTACTTGGTTGTTGCATATAAAAAACGGTAATACTATAATACTCTGTCCCTCCGGGACTAAATTACTTAGCAAGTTGACATTTGTTTATATTTAGC 2240

TTCGGGAACGTCCAACGTCTCCGGTGTAAGACCTTTCTAGTCGCATTCCTGATTCAAACAAGATGCTACTTGGTTGTATATAAAATGATTGGGCTATAAT 2340
TTCGGGAACGTCCAACGTCTCCGGTGTAAGACCTTTCTAGTCGCATTCCTGATTCAAACAAGATGCTACTTGGTTGTATATAAAATGATTGGGCTATAAT 2340

ACTCCTCTTCGCAACAATTAAATTACTCAGTAGGTGAACATTTTATATATTCAACTCTAGCATATAATACAAATTTGTTGTCGGCATAGAAATGAGTTGC 2440
ACTCCTCTTCGCAACAATTAAATTACTCAGTAGGTGAACATTTTATATATTCAACTCTAGCATATAATACAAATTTGTTGTCGGCATAGAAATGAGTTGC 2440

AATCATTTCATAAATAGGAAGAACACATGGCTTATACCAAACTAGCAGCCTAAAAAGGTTTTTTTTTCTTCTTTCTGAGAGGAGGCATTTAGCTTGTGGA 2540
AATCATTTCATAAATAGGAAGAACACATGGCTTATACCAAACTAGCAGCCTAAAAAGGTTTTTTTTTCTTCTTTCTGAGAGGAGGCATTTAGCTTGTGGA 2540
```

```
GCAAAATGTTGAAGCGGCTGGGCGAAAAAACCCTCGGCTGATCCGCGTGAGGGCACGACACGTGGCGTCCCGGTCCACGGGGTGTGTGGCCGTAGCGATT       2640
GCAAAATGTTGAAGCGGCTGGGCGAAAAAACCCTCGGCTGATCCGCGTGAGGGCACGACACGTGGCGTCCCGGTCCACGGGGTGTGTGGCCGTAGCGATT       2640

AGCGAGGCTCCGCTGCACAAAAATAGTTTACCTCTGATGCCCTTGGCCTCCCCGACGTCCCATCTCGCTTTCTCTCTTTCTCTTTCTCCCACTGGCCTGG       2740
AGCGAGGCTCCGCTGCACAAAAATAGTTTACCTCTGATGCCCTTGGCCTCCCCGACGTCCCATCTCGCTTTCTCTCTTTCTCTTTCTCCCACTGGCCTGG       2740

CCCCCTCTCCTCGTCGTCCTCCAGTCCTCATCCCCCGCCCCATGGCCACCCCCACCACGGCCGCCCCCCCCCCCCCTCCCCCCACCACTACTTCTACTCC       2840
CCCCCTCTCCTCGTCGTCCTCCAGTCCTCATCCCCCGCCCCATGGCCACCCCCACCACGGCCGCCCCCCCCCCCCCTCCCCCCACCACTACTTCTACTCC       2840

CCCCGCCCCGCCCTCGCAGCCCGCGGCCACCGCGCTCCCATGCCATAGACGCGACCCCACTCATCGGTCCAGGTC                                2915
CCCCGCCCCGCCCTCGCAGCCCGCGGCCACCGCGCTCCCATGCCATAGACGCGACCCCACTCATCGGTCCAGGTC                                2915
```

**Figure S3.** Alignment of Q gene promoter from cultivar Chinese Spring and Bobwhite. The top sequence is Chinese Spring, and the bottom sequence is cultivar Bobwhite. The promoter region is 2990 bp long upstream of the start codon of Q gene in both cultivars. The missed nucleotides in the alignment are shown as “-”; the mismatch in the alignment is shown in red fonts. The position of the last nucleotide in each row is shown on the right side of the alignment.

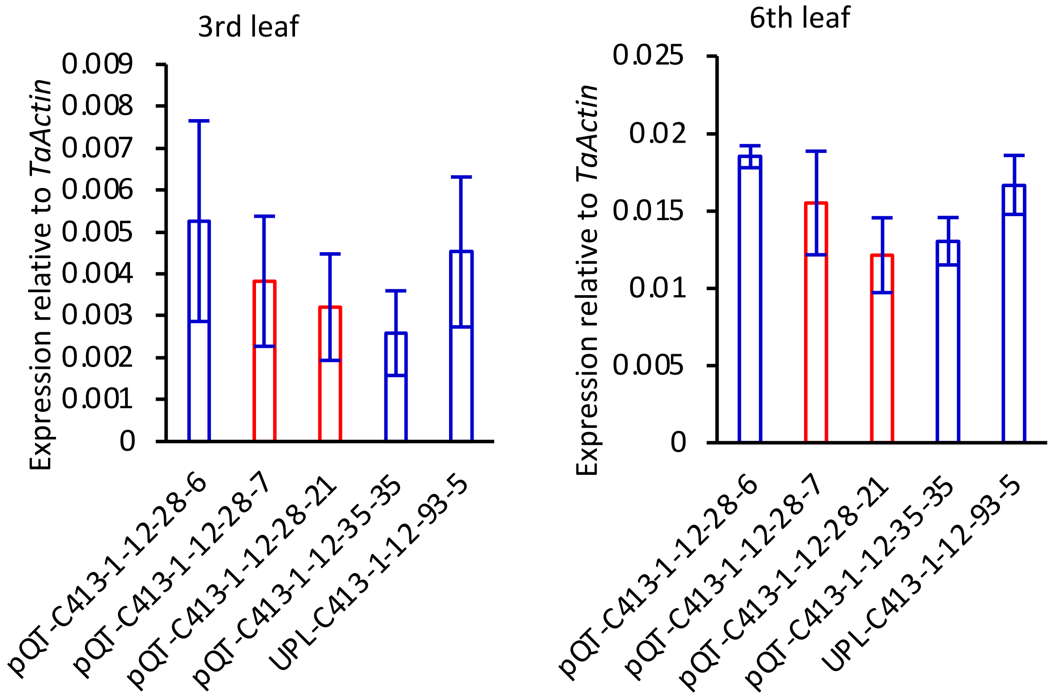


**Figure S4.** Expression of *Q* gene A genome allele in the M1 plants at 6-leaf stage. The A genome specific primers, Q5ArtF4 and Q5ArtR4 (Table S4), were applied. *TaActin* gene was used as reference. The 3rd and 6th leaf in M1 plants at 6-leaf stage were sampled for RNA isolation followed by reverse transcription to get cDNA. The plant pQT-C413-1-12-28-7 and pQT-C413-1-12-28-21 carrying homozygous long deletions are highlighted with red box in the bar plots. All the rest of the plants have wild type alleles all the promoter of Q gene. The results are shown as mean ± SE based on three technical repeats.

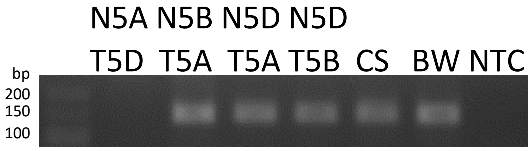


**Figure S5.** Validation of the *Q* gene A genome specific primers for RT-PCR. PCR amplification of cv. Bobwhite, cv. Chinese Spring (CS) and nullisomic-tetrasomic lines using the genome-specific primer Q5ArtF4 and Q5ArtR4 (Table S4) was performed. DNA isolated from six nullisomic-tetrasomic lines (N5A-T5D, N5B-T5A, N5DT5A, and N5DT5B), cv. Chinese Spring (CS), and cv. Bobwhite (BW). NTC is no template control of PCRs.

**Table S1.** The efficiency of editing based on the BSMV-sgRNA delivery system in transgenic wheat lines with the low (7438) and high (C413) levels of Cas9 expression.

| Plants and Constructs | Genome | Mutated Reads | Total Reads | Mutated Percentage | Average |
|---|---|---|---|---|---|
| 7438-1-3-14 BSMV-QT1 | A | 99 | 17104 | 0.58% | 0.57% |
| | B | 85 | 18766 | 0.45% | |
| | D | 128 | 18617 | 0.69% | |
| 7438-1-7-1 BSMV-QT1 | A | 145 | 43415 | 0.33% | 0.33% |
| | B | 165 | 47613 | 0.35% | |
| | D | 163 | 52476 | 0.31% | |
| 7438-1-7-2 BSMV-QT1 | A | 65 | 17375 | 0.37% | 0.38% |
| | B | 82 | 19484 | 0.42% | |
| | D | 73 | 21206 | 0.34% | |
| C413-1-12-47 BSMV-QT1 | A | 268 | 274 | 97.81% | 98.50% |
| | B | 419 | 427 | 98.13% | |
| | D | 499 | 503 | 99.20% | |
| C413-1-12-116 BSMV-QT1 | A | 381 | 385 | 98.96% | 99.03% |
| | B | 400 | 404 | 99.01% | |
| | D | 446 | 450 | 99.11% | |
| C413-1-12-117 BSMV-QT1 | A | 132 | 138 | 95.65% | 96.14% |
| | B | 162 | 164 | 98.78% | |
| | D | 204 | 216 | 94.44% | |
| C413-1-12-118 BSMV-QT1 | A | 64 | 64 | 100.00% | 99.58% |
| | B | 65 | 66 | 98.48% | |
| | D | 107 | 107 | 100.00% | |
| C413-1-12-122 BSMV-QT1 | A | 32 | 32 | 100.00% | 96.15% |
| | B | 48 | 48 | 100.00% | |
| | D | 70 | 76 | 92.11% | |

**Table S2.** The target sites selected for BSMV-sgRNA-based editing in this study.

| Gene targeted | Target name¶ | Spacer Sequence | PAM | Strand‡ targeted |
|---|---|---|---|---|
| *Q*<br>Gene ID†:<br>TraesCS5A02G473800 | QT1 | ATGAGGAACTGGACCAAGG | AGG | Plus |
| | pQT17 | GTTTCCGATGGGGCGAGCGG | GGG | Plus |
| | pQT18 | GACGTAGACCCACGCGAGGG | CGG | Minus |
| | pQT23 | CGTGCCGACGAGAGCACGGG | GGG | Minus |
| | pQT25 | AGCACAGGGCGTACGGCCGG | GGG | Plus |
| | pQT26 | CGCTACGGCCACACACCCCG | TGG | Minus |
| *TaGW7*<br>Gene ID:<br>TraesCS2A02G176000<br>TraesCS2B02G202300<br>TraesCS2D02G183400 | GW7T6 | TCCATCAACCGGGACTCGGG | AGG | Plus |
| *TaGW2*<br>Gene ID:<br>TraesCS6A02G189300<br>TraesCS6B02G215300<br>TraesCS6D02G176900 | GW2T2 | GCTCGCGCCCTGCTACCCGG | GGG | Plus |
| *TaMTL*<br>Gene ID:<br>TraesCS4A02G018100<br>TraesCS4B02G286000<br>TraesCS4D02G284700 | MTLT5 | GCATCCGCTCGCCGATCCTG | AGG | Minus |
| TaUPL3§<br>*Gene ID*:<br>TraesCS2A02G064700<br>TraesCS2B02G076900 | UPL3T11 | CAAGGAGCAGCAGGAGCCCT | CGG | Plus |

† : The gene IDs are based on the IWGSC RefSeq v1.1 gene annotation.
‡: The plus strand stand for the direction from start codon to the stop codon of a gene.
§: The Chromosome 2D genome copy of TaUPL3 is in the region of chr2D:26764500-26774638, which was not annotated. But the D genome copy is functional protein coding gene.
¶: All the CRISPR-Cas9 targets targeting the coding region of the genes except for the targets start with lower case letter p which target the promoter region of *Q* gene.

**Table S3.** Proportion of NGS reads carrying 71-bp deletion.

| M0_Plant_ID | Reads with 71 bp deletion | total reads | Ratio |
|---|---|---|---|
| C413-1-12-28 | 215 | 1947 | 11.04% |
| C413-1-12-29 | 554 | 2785 | 19.89% |
| C413-1-12-30 | 265 | 2579 | 10.28% |
| C413-1-12-31 | 393 | 2910 | 13.51% |
| C413-1-12-32 | 393 | 2580 | 15.23% |
| C413-1-12-34 | 205 | 2572 | 7.97% |
| C413-1-12-35 | 179 | 1634 | 10.95% |
| C413-1-12-36 | 321 | 2289 | 14.02% |
| C413-1-12-37 | 147 | 2841 | 5.17% |
| C413-1-12-40 | 279 | 2767 | 10.08% |

**Table S4.** The primers, oligos and synthesized double-strand DNA used in this study.

| Names | Sequences | Notes | Usage |
|---|---|---|---|
| targetVIGSsF | ATATTAATTAACGCTTGCTGCATCAGACTTG | Pac1 cut site is in red | Amplify sgRNA from CRISPR-Cas9 plasmids and insert into BSMV gamma chain plasmid on the sense direction. |
| targetVIGSsR | TATGCGGCCGCAAAGCACCGACTCGGTGCC | Not1 cut site is in red | |
| targetVIGSaF | ATATTAATTAAAAAGCACCGACTCGGTGCC | Pac1 cut site is in red | Amplify sgRNA from CRISPR-Cas9 plasmids and insert into BSMV gamma chain plasmid on the antisense direction. |
| targetVIGSaR | TATGCGGCCGCCGCTTGCTGCATCAGACTTG | Not1 cut site is in red | |
| VIGS-GW2T2sgRNA-F | CTTCCGTTGCTAGCTGATTAATTAAgctcgcgccctgctacccggGTTTTAGAGCTAGAA | Capital letters on the 5' end overlap with PacI and NotI digested pBSMVgammaPDS, lower case letters in the middle are the spacer of GW2T2, capital letters on the 3' end is specific primers fo 5' end of sgRNA scafold. | Amplify sgRNA from CRISPR-Cas9 plasmids and insert into BSMV gamma chain plasmid using Gibson reaction. |
| VIGS-sgRNA-R | tttttttagctagctgagcggccgcGCACCGACTCGGTGCCACTT | Lower case letters on the 5' end overlap with PacI and NotI digested pBSMVgammaPDS, capital letters on the 3' end is specific primers fo 3' end of sgRNA scafold. | |
| SgRNA scalfold::AtFT_CDS | CTTCCGTTGCTAGCTGATTAATTAAgctcgcgccctgctacccggGTTTTAGAGCTAGAAATAGCAAGTTAAAATAAGGCTAGTCCGTTATCAACTTGAAAAAGTGGCACCGAGTCGGTGCatgtctataaatataagagaccctcttatagtaagcagagttgttggagacgttcttgatccgtttaatagatcaatcactctaaaggttacttatggccaaagagaggtgactaatggcttggatctaaggccttctcaggttcaaaacaagccaagagttgagattggtggagaagacctcaggaacttctatactttggttatggtggatccagatgttccaagtcctagcaaccctcacctccgagaatatctccattggttggtgactgatatccctgctacaactggaacaacctttggcaatgagattgtgtgttacgaaaatccaagtcccactgcaggaattcatcgtgtcgtgtttatattgtttcgacagcttggcaggcaaacagtgtatgcaccagggtggcgccagaacttcaacactcgcgagtttgctgagatctacaatctcggccttcccgtggccgcagttttctacaattgtcagagggagagtggctgcggaggaagaagactttagGCGGCCGCTCAGCTAGCTAAAAAAA | Each part are differenttiated with upper and lower case letters. From left to right, overlap with plasmid, spacer, sgRNA scalfold, mobile elements, overlap with plasmid. | Construct BSMV gamma chain plasmid with fused sgRNA and mobile elements. |
| SgRNA scalfold::Vrn3 CDS | CTTCCGTTGCTAGCTGATTAATTAAgctcgcgccctgctacccggGTTTTAGAGCTAGAAATAGCAAGTTAAAATAAGGCTAGTCCGTTATCAACTTGAAAAAGTGGCACCGAGTCGGTGCatggccggtagggatagggacccgctggtggttggcagggttgtgggggacgtgctggaccccttcgtccggaccaccaacctcagggtgaccttcgggaacaggaccgtgtccaacggctgcgagctcaagccgtccatggtcgcccagcagcccagggttgaggtgggcggcaatgagatgaggaccttctacacactcgtgatggtagacccagatgctccaagtccaagcgatcccaaccttagggagtatctccactggcttgtgacagatatccccggtacaactggtgcgtcgttcgggcaggaggtgatgtgctacgagagccctcgtccgaccatggggatccaccgcttcgtgctcgtactcttccagcagctcgggcggcagacggtgtacgcccccgggtggcgccagaacttcaacaccagggacttcgccgagctctacaacctcggcccgcctgtcgccgccgtctacttcaactgccagcgtgaggccggctccggcggcaggaggatgtacaattgaGCGGCCGCTCAGCTAGCTAAAAAAA | | |

| | | | |
|---|---|---|---|
| SgRNA scalfold::tRNA$^{Met}$ | CTTCCGTTGCTAGCTGATTAATTAAgctcgcgccctgctacccggGTTTTAGAGCTAGAAATAGCAAGTTAAAATAAGGCTAGTCCGTTATCAACTTGAAAAAGTGGCACCGAGTCGGTGCatcagagtggcgcagcggaagcgtggtgggcccataacccacaggtcccaggatcgaaacctggctctgataGCGGCCGCTCAGCTAGCTAAAAAAA | | |
| SgRNA scalfold::tRNA$^{Ile}$ | CTTCCGTTGCTAGCTGATTAATTAAgctcgcgccctgctacccggGTTTTAGAGCTAGAAATAGCAAGTTAAAATAAGGCTAGTCCGTTATCAACTTGAAAAAGTGGCACCGAGTCGGTGCgctcccgtagctcagttggttagagcgttggtcttatgagccgaaggtcgcgggttcgagccccgccgggagcaGCGGCCGCTCAGCTAGCTAAAAAAA | | |
| BSMV-UPL3T11-SgRNA | CTTCCGTTGCTAGCTGATTAATTAAcaaggagcagcaggagccctGTTTTAGAGCTAGAAATAGCAAGTTAAAATAAGGCTAGTCCGTTATCAACTTGAAAAAGTGGCACCGAGTCGGTGCgcggccgctcagctagctaaaaaaa | Each part are differenttiated with upper and lower case letters. From left to right, overlap with plasmid, spacer, sgRNA scalfold, overlap with plasmid. | Construct BSMV gamma chain plasmid by inserting into BSMV gamma chain plasmid using Gibson reaction. |
| VIGS-pQT17FsgRNA-F | CTTCCGTTGCTAGCTGATTAATTAAgtttccgatggggcgagcggGTTTTAGAGCTAGAA | Capital letters on the 5' end overlap with PacI and NotI digested pBSMVgammaPDS, lower case letters in the middle are the spacer of target sites, capital letters on the 3' end is specific primers fo 5' end of sgRNA scafold. | Work in pair with primer VIGS-sgRNA-R, amplify sgRNA from CRISPR-Cas9 plasmids and insert into BSMV gamma chain plasmid using Gibson reaction. |
| VIGS-pQT18FsgRNA-F | CTTCCGTTGCTAGCTGATTAATTAAgacgtagacccacgcgagggGTTTTAGAGCTAGAA | | |
| VIGS-pQT23FsgRNA-F | CTTCCGTTGCTAGCTGATTAATTAAcgtgccgacgagagcacgggGTTTTAGAGCTAGAA | | |
| VIGS-pQT25FsgRNA-F | CTTCCGTTGCTAGCTGATTAATTAAagcacagggcgtacggccggGTTTTAGAGCTAGAA | | |
| VIGS-pQT26FsgRNA-F | CTTCCGTTGCTAGCTGATTAATTAAcgctacggccacacaccccgGTTTTAGAGCTAGAA | | |
| VIGS-GW7T6FsgRNA-F | CTTCCGTTGCTAGCTGATTAATTAAtccatcaaccgggactcgggGTTTTAGAGCTAGAA | | |
| BSMVseqL | GGAGCTGAAACTTTCTCATATCG | | Sanger sequencing of pBSMVgamma constructs with insertions |
| BSMVseqR | GTGGACTTGCAAACACTCCC | | |
| Q5ArtF4 | CTGATGCTCTTGACTTGGATCTG | | Check the expression of Q gene A genome allele |
| Q5ArtR4 | TTTTGTTTCTTTACCTGAGAAGAGAT | | |
| zCAS9-F | CGGACTAGTATGGATTACAAGGACCACGACG | | Check the expression of Cas9 in transgenic plants |
| zCAS9seq1 | accttgtactcgtcggtgatcac | | |
| TaActin-F | ACCTTCAGTTGCCCAGCAAT | | Amplify TaActin gene as reference to check the expression level of other genes. |
| TaActin-R | CAGAGTCGAGCACAATACCAGTTG | | |
| QT1MiseqF1 | ctctttccctacacgacgctcttccgatctCGCTTtggtttgtccgatggttgat | The target specific primers are shown as lower case letter on the 3' end, the 5 additional barcoding nucleotides are shown as upper case letters, the 5' end lower case letters are part of Illumina Truseq adapter. | The first round PCR primers for NGS library of target site QT1. |
| QT1MiseqF2 | ctctttccctacacgacgctcttccgatctCTAGCtggtttgtccgatggttgat | | |
| QT1MiseqF3 | ctctttccctacacgacgctcttccgatctACAAAtggtttgtccgatggttgat | | |
| QT1MiseqF4 | ctctttccctacacgacgctcttccgatctTTCTCtggtttgtccgatggttgat | | |
| QT1MiseqF5 | ctctttccctacacgacgctcttccgatctAGCCCtggtttgtccgatggttgat | | |
| QT1MiseqF6 | ctctttccctacacgacgctcttccgatctGTATTtggtttgtccgatggttgat | | |
| QT1MiseqF7 | ctctttccctacacgacgctcttccgatctCTGTAtggtttgtccgatggttgat | | |
| QT1MiseqF8 | ctctttccctacacgacgctcttccgatctACCGTtggtttgtccgatggttgat | | |
| QT1MiseqR1 | ctggagttcagacgtgtgctcttccgatctGCTTAtgtgcctgcttacttcttgc | | |
| QT1MiseqR2 | ctggagttcagacgtgtgctcttccgatctGGTGTtgtgcctgcttacttcttgc | | |
| QT1MiseqR3 | ctggagttcagacgtgtgctcttccgatctAGGATtgtgcctgcttacttcttgc | | |

| | | | |
|---|---|---|---|
| QT1MiseqR4 | ctggagttcagacgtgtgctcttccgatctATTGAtgtgcctgcttacttcttgc | | |
| QT1MiseqR5 | ctggagttcagacgtgtgctcttccgatctCATCTtgtgcctgcttacttcttgc | | |
| QT1MiseqR6 | ctggagttcagacgtgtgctcttccgatctCCTACtgtgcctgcttacttcttgc | | |
| QT1MiseqR7 | ctggagttcagacgtgtgctcttccgatctGAGGAtgtgcctgcttacttcttgc | | |
| QT1MiseqR8 | ctggagttcagacgtgtgctcttccgatctGGAACtgtgcctgcttacttcttgc | | |
| QT1MiseqR9 | ctggagttcagacgtgtgctcttccgatctGTCAAtgtgcctgcttacttcttgc | | |
| QT1MiseqR10 | ctggagttcagacgtgtgctcttccgatctTAATAtgtgcctgcttacttcttgc | | |
| QT1MiseqR11 | ctggagttcagacgtgtgctcttccgatctTACATtgtgcctgcttacttcttgc | | |
| QT1MiseqR12 | ctggagttcagacgtgtgctcttccgatctTCGTTtgtgcctgcttacttcttgc | | |
| GW2T2MiseqF1 | ctctttccctacacgacgctcttccgatctCTCCatggggaacagaataggagg | The target specific primers are shown as lower case letter on the 3' end, the 5 additional barcoding nucleotides are shown as upper case letters, the 5' end lower case letters are part of Illumina Truseq adapter. | The first round PCR primers for NGS library of target site GW2T2. |
| GW2T2MiseqF2 | ctctttccctacacgacgctcttccgatctTGCAatggggaacagaataggagg | | |
| GW2T2MiseqF3 | ctctttccctacacgacgctcttccgatctACTAatggggaacagaataggagg | | |
| GW2T2MiseqF4 | ctctttccctacacgacgctcttccgatctCAGAatggggaacagaataggagg | | |
| GW2T2MiseqF5 | ctctttccctacacgacgctcttccgatctAACTatggggaacagaataggagg | | |
| GW2T2MiseqF6 | ctctttccctacacgacgctcttccgatctGCGTatggggaacagaataggagg | | |
| GW2T2MiseqF7 | ctctttccctacacgacgctcttccgatctCGATatggggaacagaataggagg | | |
| GW2T2MiseqF8 | ctctttccctacacgacgctcttccgatctGTAAatggggaacagaataggagg | | |
| GW2T2MiseqR1 | ctggagttcagacgtgtgctcttccgatctCTCCaggaagcagatggggcactc | | |
| GW2T2MiseqR2 | ctggagttcagacgtgtgctcttccgatctTGCAaggaagcagatggggcactc | | |
| GW2T2MiseqR3 | ctggagttcagacgtgtgctcttccgatctACTAaggaagcagatggggcactc | | |
| GW2T2MiseqR4 | ctggagttcagacgtgtgctcttccgatctCAGAaggaagcagatggggcactc | | |
| GW2T2MiseqR5 | ctggagttcagacgtgtgctcttccgatctAACTaggaagcagatggggcactc | | |
| GW2T2MiseqR6 | ctggagttcagacgtgtgctcttccgatctGCGTaggaagcagatggggcactc | | |
| GW2T2MiseqR7 | ctggagttcagacgtgtgctcttccgatctCGATaggaagcagatggggcactc | | |
| GW2T2MiseqR8 | ctggagttcagacgtgtgctcttccgatctGTAAaggaagcagatggggcactc | | |
| GW2T2MiseqR9 | ctggagttcagacgtgtgctcttccgatctAGGCaggaagcagatggggcactc | | |
| GW2T2MiseqR10 | ctggagttcagacgtgtgctcttccgatctGATCaggaagcagatggggcactc | | |
| GW2T2MiseqR11 | ctggagttcagacgtgtgctcttccgatctTCACaggaagcagatggggcactc | | |
| GW2T2MiseqR12 | ctggagttcagacgtgtgctcttccgatctTGCGAaggaagcagatggggcactc | | |
| GW7T6MiseqF1 | ctctttccctacacgacgctcttccgatctCGCTTgacctttcggtttattttgcag | The target specific primers are shown as lower case letter on the 3' end, the 5 additional barcoding nucleotides are shown as upper case letters, the 5' end lower case letters are part of Illumina Truseq adapter. | The first round PCR primers for NGS library of target site GW7T6. |
| GW7T6MiseqF2 | ctctttccctacacgacgctcttccgatctCTAGCgacctttcggtttattttgcag | | |
| GW7T6MiseqF3 | ctctttccctacacgacgctcttccgatctACAAAgacctttcggtttattttgcag | | |
| GW7T6MiseqF4 | ctctttccctacacgacgctcttccgatctTTCTCgacctttcggtttattttgcag | | |
| GW7T6MiseqF5 | ctctttccctacacgacgctcttccgatctAGCCCgacctttcggtttattttgcag | | |

| | | | |
|---|---|---|---|
| GW7T6MiseqF6 | ctctttccctacacgacgctcttccgatctGTATTgac ctttcggtttattttgcag | | |
| GW7T6MiseqF7 | ctctttccctacacgacgctcttccgatctCTGTAgac ctttcggtttattttgcag | | |
| GW7T6MiseqF8 | ctctttccctacacgacgctcttccgatctACCGTgac ctttcggtttattttgcag | | |
| GW7T6MiseqR1 | ctggagttcagacgtgtgctcttccgatctGCTTAgta ggttccgtccatcgattt | | |
| GW7T6MiseqR2 | ctggagttcagacgtgtgctcttccgatctGGTGTgta ggttccgtccatcgattt | | |
| GW7T6MiseqR3 | ctggagttcagacgtgtgctcttccgatctAGGATgta ggttccgtccatcgattt | | |
| GW7T6MiseqR4 | ctggagttcagacgtgtgctcttccgatctATTGAgta ggttccgtccatcgattt | | |
| GW7T6MiseqR5 | ctggagttcagacgtgtgctcttccgatctCATCTgta ggttccgtccatcgattt | | |
| GW7T6MiseqR6 | ctggagttcagacgtgtgctcttccgatctCCTACgta ggttccgtccatcgattt | | |
| GW7T6MiseqR7 | ctggagttcagacgtgtgctcttccgatctGAGGAgta ggttccgtccatcgattt | | |
| GW7T6MiseqR8 | ctggagttcagacgtgtgctcttccgatctGGAACgta ggttccgtccatcgattt | | |
| GW7T6MiseqR9 | ctggagttcagacgtgtgctcttccgatctGTCAAgta ggttccgtccatcgattt | | |
| GW7T6MiseqR10 | ctggagttcagacgtgtgctcttccgatctTAATAgta ggttccgtccatcgattt | | |
| GW7T6MiseqR11 | ctggagttcagacgtgtgctcttccgatctTACATgta ggttccgtccatcgattt | | |
| GW7T6MiseqR12 | ctggagttcagacgtgtgctcttccgatctTCGTTgta ggttccgtccatcgattt | | |
| UPL3T11MiF1 | ctctttccctacacgacgctcttccgatctCGCTTccc aaccctaaccctagcc | The target specific primers are shown as lower case letter on the 3' end, the 5 additional barcoding nucleotides are shown as upper case letters, the 5' end lower case letters are part of Illumina Truseq adapter. | The first round PCR primers for NGS library of target site UPL3T11. |
| UPL3T11MiF2 | ctctttccctacacgacgctcttccgatctCTAGCccc aaccctaaccctagcc | | |
| UPL3T11MiF3 | ctctttccctacacgacgctcttccgatctACAAAccc aaccctaaccctagcc | | |
| UPL3T11MiF4 | ctctttccctacacgacgctcttccgatctTTCTCccc aaccctaaccctagcc | | |
| UPL3T11MiF5 | ctctttccctacacgacgctcttccgatctAGCCCccc aaccctaaccctagcc | | |
| UPL3T11MiF6 | ctctttccctacacgacgctcttccgatctGTATTccc aaccctaaccctagcc | | |
| UPL3T11MiF7 | ctctttccctacacgacgctcttccgatctCTGTAccc aaccctaaccctagcc | | |
| UPL3T11MiF8 | ctctttccctacacgacgctcttccgatctACCGTccc aaccctaaccctagcc | | |
| UPL3T11MiR1 | ctggagttcagacgtgtgctcttccgatctGCTTActc gtcgtcgtcgtccat | | |
| UPL3T11MiR2 | ctggagttcagacgtgtgctcttccgatctGGTGTctc gtcgtcgtcgtccat | | |
| UPL3T11MiR3 | ctggagttcagacgtgtgctcttccgatctAGGATctc gtcgtcgtcgtccat | | |
| UPL3T11MiR4 | ctggagttcagacgtgtgctcttccgatctATTGActc gtcgtcgtcgtccat | | |
| UPL3T11MiR5 | ctggagttcagacgtgtgctcttccgatctCATCTctc gtcgtcgtcgtccat | | |
| UPL3T11MiR6 | ctggagttcagacgtgtgctcttccgatctCCTACctc gtcgtcgtcgtccat | | |
| UPL3T11MiR7 | ctggagttcagacgtgtgctcttccgatctGAGGActc gtcgtcgtcgtccat | | |
| UPL3T11MiR8 | ctggagttcagacgtgtgctcttccgatctGGAACctc gtcgtcgtcgtccat | | |
| UPL3T11MiR9 | ctggagttcagacgtgtgctcttccgatctGTCAActc gtcgtcgtcgtccat | | |
| UPL3T11MiR10 | ctggagttcagacgtgtgctcttccgatctTAATActc gtcgtcgtcgtccat | | |
| UPL3T11MiR11 | ctggagttcagacgtgtgctcttccgatctTACATctc gtcgtcgtcgtccat | | |

| | | | |
|---|---|---|---|
| UPL3T11MiR12 | ctggagttcagacgtgtgctcttccgatctTCGTTctc gtcgtcgtcgtccat | | |
| pQT17MiF1 | ctctttccctacacgacgctcttccgatctCGCTTgtg accacgagcgtaaggtt | The target specific primers are shown as lower case letter on the 3' end, the 5 additional barcoding nucleotides are shown as upper case letters, the 5' end lower case letters are part of Illumina Truseq adapter. | The first round PCR primers for NGS library of target site pQT17. |
| pQT17MiF2 | ctctttccctacacgacgctcttccgatctCTAGCgtg accacgagcgtaaggtt | | |
| pQT17MiF3 | ctctttccctacacgacgctcttccgatctACAAAgtg accacgagcgtaaggtt | | |
| pQT17MiF4 | ctctttccctacacgacgctcttccgatctTTCTCgtg accacgagcgtaaggtt | | |
| pQT17MiF5 | ctctttccctacacgacgctcttccgatctAGCCCgtg accacgagcgtaaggtt | | |
| pQT17MiF6 | ctctttccctacacgacgctcttccgatctGTATTgtg accacgagcgtaaggtt | | |
| pQT17MiF7 | ctctttccctacacgacgctcttccgatctCTGTAgtg accacgagcgtaaggtt | | |
| pQT17MiF8 | ctctttccctacacgacgctcttccgatctACCGTgtg accacgagcgtaaggtt | | |
| pQT17MiR1 | ctggagttcagacgtgtgctcttccgatctGCTTAcca caggcgtacctgtcc | | |
| pQT17MiR2 | ctggagttcagacgtgtgctcttccgatctGGTGTcca caggcgtacctgtcc | | |
| pQT17MiR3 | ctggagttcagacgtgtgctcttccgatctAGGATcca caggcgtacctgtcc | | |
| pQT17MiR4 | ctggagttcagacgtgtgctcttccgatctATTGAcca caggcgtacctgtcc | | |
| pQT17MiR5 | ctggagttcagacgtgtgctcttccgatctCATCTcca caggcgtacctgtcc | | |
| pQT17MiR6 | ctggagttcagacgtgtgctcttccgatctCCTACcca caggcgtacctgtcc | | |
| pQT17MiR7 | ctggagttcagacgtgtgctcttccgatctGAGGAcca caggcgtacctgtcc | | |
| pQT17MiR8 | ctggagttcagacgtgtgctcttccgatctGGAACcca caggcgtacctgtcc | | |
| pQT17MiR9 | ctggagttcagacgtgtgctcttccgatctGTCAAcca caggcgtacctgtcc | | |
| pQT17MiR10 | ctggagttcagacgtgtgctcttccgatctTAATAcca caggcgtacctgtcc | | |
| pQT17MiR11 | ctggagttcagacgtgtgctcttccgatctTACATcca caggcgtacctgtcc | | |
| pQT17MiR12 | ctggagttcagacgtgtgctcttccgatctTCGTTcca caggcgtacctgtcc | | |
| pQT18MiF1 | ctctttccctacacgacgctcttccgatctCGCTTaag gaacaacgtttccgatg | The target specific primers are shown as lower case letter on the 3' end, the 5 additional barcoding nucleotides are shown as upper case letters, the 5' end lower case letters are part of Illumina Truseq adapter. | The first round PCR primers for NGS library of target site pQT18. |
| pQT18MiF2 | ctctttccctacacgacgctcttccgatctCTAGCaag gaacaacgtttccgatg | | |
| pQT18MiF3 | ctctttccctacacgacgctcttccgatctACAAAaag gaacaacgtttccgatg | | |
| pQT18MiF4 | ctctttccctacacgacgctcttccgatctTTCTCaag gaacaacgtttccgatg | | |
| pQT18MiF5 | ctctttccctacacgacgctcttccgatctAGCCCaag gaacaacgtttccgatg | | |
| pQT18MiF6 | ctctttccctacacgacgctcttccgatctGTATTaag gaacaacgtttccgatg | | |
| pQT18MiF7 | ctctttccctacacgacgctcttccgatctCTGTAaag gaacaacgtttccgatg | | |
| pQT18MiF8 | ctctttccctacacgacgctcttccgatctACCGTaag gaacaacgtttccgatg | | |
| pQT18MiR1 | ctggagttcagacgtgtgctcttccgatctGCTTAcgc tctctttcgtgtgtgtg | | |
| pQT18MiR2 | ctggagttcagacgtgtgctcttccgatctGGTGTcgc tctctttcgtgtgtgtg | | |
| pQT18MiR3 | ctggagttcagacgtgtgctcttccgatctAGGATcgc tctctttcgtgtgtgtg | | |
| pQT18MiR4 | ctggagttcagacgtgtgctcttccgatctATTGAcgc tctctttcgtgtgtgtg | | |
| pQT18MiR5 | ctggagttcagacgtgtgctcttccgatctCATCTcgc tctctttcgtgtgtgtg | | |

| | | | |
|---|---|---|---|
| pQT18MiR6 | ctggagttcagacgtgtgctcttccgatctCCTACcgc tctctttcgtgtgtgtg | | |
| pQT18MiR7 | ctggagttcagacgtgtgctcttccgatctGAGGAcgc tctctttcgtgtgtgtg | | |
| pQT18MiR8 | ctggagttcagacgtgtgctcttccgatctGGAACcgc tctctttcgtgtgtgtg | | |
| pQT18MiR9 | ctggagttcagacgtgtgctcttccgatctGTCAAcgc tctctttcgtgtgtgtg | | |
| pQT18MiR10 | ctggagttcagacgtgtgctcttccgatctTAATAcgc tctctttcgtgtgtgtg | | |
| pQT18MiR11 | ctggagttcagacgtgtgctcttccgatctTACATcgc tctctttcgtgtgtgtg | | |
| pQT18MiR12 | ctggagttcagacgtgtgctcttccgatctTCGTTcgc tctctttcgtgtgtgtg | | |
| pQT23MiF1 | ctctttccctacacgacgctcttccgatctCGCTTaat tgcacccatcccattac | The target specific primers are shown as lower case letter on the 3' end, the 5 additional barcoding nucleotides are shown as upper case letters, the 5' end lower case letters are part of Illumina Truseq adapter. | The first round PCR primers for NGS library of target site pQT23. |
| pQT23MiF2 | ctctttccctacacgacgctcttccgatctCTAGCaat tgcacccatcccattac | | |
| pQT23MiF3 | ctctttccctacacgacgctcttccgatctACAAAaat tgcacccatcccattac | | |
| pQT23MiF4 | ctctttccctacacgacgctcttccgatctTTCTCaat tgcacccatcccattac | | |
| pQT23MiF5 | ctctttccctacacgacgctcttccgatctAGCCCaat tgcacccatcccattac | | |
| pQT23MiF6 | ctctttccctacacgacgctcttccgatctGTATTaat tgcacccatcccattac | | |
| pQT23MiF7 | ctctttccctacacgacgctcttccgatctCTGTAaat tgcacccatcccattac | | |
| pQT23MiF8 | ctctttccctacacgacgctcttccgatctACCGTaat tgcacccatcccattac | | |
| pQT23MiR1 | ctggagttcagacgtgtgctcttccgatctGCTTAagg cctcgtgtgttgacc | | |
| pQT23MiR2 | ctggagttcagacgtgtgctcttccgatctGGTGTagg cctcgtgtgttgacc | | |
| pQT23MiR3 | ctggagttcagacgtgtgctcttccgatctAGGATagg cctcgtgtgttgacc | | |
| pQT23MiR4 | ctggagttcagacgtgtgctcttccgatctATTGAagg cctcgtgtgttgacc | | |
| pQT23MiR5 | ctggagttcagacgtgtgctcttccgatctCATCTagg cctcgtgtgttgacc | | |
| pQT23MiR6 | ctggagttcagacgtgtgctcttccgatctCCTACagg cctcgtgtgttgacc | | |
| pQT23MiR7 | ctggagttcagacgtgtgctcttccgatctGAGGAagg cctcgtgtgttgacc | | |
| pQT23MiR8 | ctggagttcagacgtgtgctcttccgatctGGAACagg cctcgtgtgttgacc | | |
| pQT23MiR9 | ctggagttcagacgtgtgctcttccgatctGTCAAagg cctcgtgtgttgacc | | |
| pQT23MiR10 | ctggagttcagacgtgtgctcttccgatctTAATAagg cctcgtgtgttgacc | | |
| pQT23MiR11 | ctggagttcagacgtgtgctcttccgatctTACATagg cctcgtgtgttgacc | | |
| pQT23MiR12 | ctggagttcagacgtgtgctcttccgatctTCGTTagg cctcgtgtgttgacc | | |
| pQT25MiF1 | ctctttccctacacgacgctcttccgatctCGCTTtgc acttgggcaatgtaatg | The target specific primers are shown as lower case letter on the 3' end, the 5 additional barcoding nucleotides are shown as upper case letters, the 5' end lower case letters are part of Illumina Truseq adapter. | The first round PCR primers for NGS library of target site pQT25. |
| pQT25MiF2 | ctctttccctacacgacgctcttccgatctCTAGCtgc acttgggcaatgtaatg | | |
| pQT25MiF3 | ctctttccctacacgacgctcttccgatctACAAAtgc acttgggcaatgtaatg | | |
| pQT25MiF4 | ctctttccctacacgacgctcttccgatctTTCTCtgc acttgggcaatgtaatg | | |
| pQT25MiF5 | ctctttccctacacgacgctcttccgatctAGCCCtgc acttgggcaatgtaatg | | |
| pQT25MiF6 | ctctttccctacacgacgctcttccgatctGTATTtgc acttgggcaatgtaatg | | |
| pQT25MiF7 | ctctttccctacacgacgctcttccgatctCTGTAtgc acttgggcaatgtaatg | | |

| | | | |
|---|---|---|---|
| pQT25MiF8 | ctctttccctacacgacgctcttccgatctACCGTtgcacttgggcaatgtaatg | | |
| pQT25MiR1 | ctggagttcagacgtgtgctcttccgatctGCTTAagtggcatggacaaagaacc | | |
| pQT25MiR2 | ctggagttcagacgtgtgctcttccgatctGGTGTagtggcatggacaaagaacc | | |
| pQT25MiR3 | ctggagttcagacgtgtgctcttccgatctAGGATagtggcatggacaaagaacc | | |
| pQT25MiR4 | ctggagttcagacgtgtgctcttccgatctATTGAagtggcatggacaaagaacc | | |
| pQT25MiR5 | ctggagttcagacgtgtgctcttccgatctCATCTagtggcatggacaaagaacc | | |
| pQT25MiR6 | ctggagttcagacgtgtgctcttccgatctCCTACagtggcatggacaaagaacc | | |
| pQT25MiR7 | ctggagttcagacgtgtgctcttccgatctGAGGAagtggcatggacaaagaacc | | |
| pQT25MiR8 | ctggagttcagacgtgtgctcttccgatctGGAACagtggcatggacaaagaacc | | |
| pQT25MiR9 | ctggagttcagacgtgtgctcttccgatctGTCAAagtggcatggacaaagaacc | | |
| pQT25MiR10 | ctggagttcagacgtgtgctcttccgatctTAATAagtggcatggacaaagaacc | | |
| pQT25MiR11 | ctggagttcagacgtgtgctcttccgatctTACATagtggcatggacaaagaacc | | |
| pQT25MiR12 | ctggagttcagacgtgtgctcttccgatctTCGTTagtggcatggacaaagaacc | | |
| pQT26MiF1 | ctctttccctacacgacgctcttccgatctCGCTTaggaggcatttagcttgtggag | The target specific primers are shown as lower case letter on the 3' end, the 5 additional barcoding nucleotides are shown as upper case letters, the 5' end lower case letters are part of Illumina Truseq adapter. | The first round PCR primers for NGS library of target site pQT26. |
| pQT26MiF2 | ctctttccctacacgacgctcttccgatctCTAGCaggaggcatttagcttgtggag | | |
| pQT26MiF3 | ctctttccctacacgacgctcttccgatctACAAAaggaggcatttagcttgtggag | | |
| pQT26MiF4 | ctctttccctacacgacgctcttccgatctTTCTCaggaggcatttagcttgtggag | | |
| pQT26MiF5 | ctctttccctacacgacgctcttccgatctAGCCCaggaggcatttagcttgtggag | | |
| pQT26MiF6 | ctctttccctacacgacgctcttccgatctGTATTaggaggcatttagcttgtggag | | |
| pQT26MiF7 | ctctttccctacacgacgctcttccgatctCTGTAaggaggcatttagcttgtggag | | |
| pQT26MiF8 | ctctttccctacacgacgctcttccgatctACCGTaggaggcatttagcttgtggag | | |
| pQT26MiR1 | ctggagttcagacgtgtgctcttccgatctGCTTAgccagtgggagaaagagaaaga | | |
| pQT26MiR2 | ctggagttcagacgtgtgctcttccgatctGGTGTgccagtgggagaaagagaaaga | | |
| pQT26MiR3 | ctggagttcagacgtgtgctcttccgatctAGGATgccagtgggagaaagagaaaga | | |
| pQT26MiR4 | ctggagttcagacgtgtgctcttccgatctATTGAgccagtgggagaaagagaaaga | | |
| pQT26MiR5 | ctggagttcagacgtgtgctcttccgatctCATCTgccagtgggagaaagagaaaga | | |
| pQT26MiR6 | ctggagttcagacgtgtgctcttccgatctCCTACgccagtgggagaaagagaaaga | | |
| pQT26MiR7 | ctggagttcagacgtgtgctcttccgatctGAGGAgccagtgggagaaagagaaaga | | |
| pQT26MiR8 | ctggagttcagacgtgtgctcttccgatctGGAACgccagtgggagaaagagaaaga | | |
| pQT26MiR9 | ctggagttcagacgtgtgctcttccgatctGTCAAgccagtgggagaaagagaaaga | | |
| pQT26MiR10 | ctggagttcagacgtgtgctcttccgatctTAATAgccagtgggagaaagagaaaga | | |
| pQT26MiR11 | ctggagttcagacgtgtgctcttccgatctTACATgccagtgggagaaagagaaaga | | |
| pQT26MiR12 | ctggagttcagacgtgtgctcttccgatctTCGTTgccagtgggagaaagagaaaga | | |
| PCR_Truseq_Amp_F | AATGATACGGCGACCACCGAGATCTACACTCTTTCCCTACACGAC | | |

| | | | |
|---|---|---|---|
| PCR_Truseq_Amp_R_1 | CAAGCAGAAGACGGCATACGAGATCGTGATGTGACTGGAGTTCAGACG | The red color upper case letters are Illumina Truseq barcodes, the black color upper case letters are part of Illumina Truseq adapter. PCR_Truseq_Amp_F is the forward primer while all others are reverse primers. The numbers in the name of reverse primers are the number of Truseq barcodes. | The second round PCR for NGS libraries. |
| PCR_Truseq_Amp_R_2 | CAAGCAGAAGACGGCATACGAGATACATCGGTGACTGGAGTTCAGACG | | |
| PCR_Truseq_Amp_R_3 | CAAGCAGAAGACGGCATACGAGATGCCTAAGTGACTGGAGTTCAGACG | | |
| PCR_Truseq_Amp_R_4 | CAAGCAGAAGACGGCATACGAGATTGGTCAGTGACTGGAGTTCAGACG | | |
| PCR_Truseq_Amp_R_5 | CAAGCAGAAGACGGCATACGAGATCACTGTGTGACTGGAGTTCAGACG | | |
| PCR_Truseq_Amp_R_6 | CAAGCAGAAGACGGCATACGAGATATTGGCGTGACTGGAGTTCAGACG | | |
| PCR_Truseq_Amp_R_7 | CAAGCAGAAGACGGCATACGAGATGATCTGGTGACTGGAGTTCAGACG | | |
| PCR_Truseq_Amp_R_8 | CAAGCAGAAGACGGCATACGAGATTCAAGTGTGACTGGAGTTCAGACG | | |
| PCR_Truseq_Amp_R_9 | CAAGCAGAAGACGGCATACGAGATCTGATCGTGACTGGAGTTCAGACG | | |
| PCR_Truseq_Amp_R_10 | CAAGCAGAAGACGGCATACGAGATAAGCTAGTGACTGGAGTTCAGACG | | |
| PCR_Truseq_Amp_R_11 | CAAGCAGAAGACGGCATACGAGATGTAGCCGTGACTGGAGTTCAGACG | | |
| PCR_Truseq_Amp_R_12 | CAAGCAGAAGACGGCATACGAGATTACAAGGTGACTGGAGTTCAGACG | | |
| PCR_Truseq_Amp_R_13 | CAAGCAGAAGACGGCATACGAGATTTGACTGTGACTGGAGTTCAGACG | | |
| PCR_Truseq_Amp_R_14 | CAAGCAGAAGACGGCATACGAGATGGAACTGTGACTGGAGTTCAGACG | | |
| PCR_Truseq_Amp_R_15 | CAAGCAGAAGACGGCATACGAGATTGACATGTGACTGGAGTTCAGACG | | |
| PCR_Truseq_Amp_R_16 | CAAGCAGAAGACGGCATACGAGATGGACGGGTGACTGGAGTTCAGACG | | |
| PCR_Truseq_Amp_R_18 | CAAGCAGAAGACGGCATACGAGATGCGGACGTGACTGGAGTTCAGACG | | |
| PCR_Truseq_Amp_R_19 | CAAGCAGAAGACGGCATACGAGATTTTCACGTGACTGGAGTTCAGACG | | |
| PCR_Truseq_Amp_R_20 | CAAGCAGAAGACGGCATACGAGATGGCCACGTGACTGGAGTTCAGACG | | |
| PCR_Truseq_Amp_R_21 | CAAGCAGAAGACGGCATACGAGATCGAAACGTGACTGGAGTTCAGACG | | |
| PCR_Truseq_Amp_R_22 | CAAGCAGAAGACGGCATACGAGATCGTACGGTGACTGGAGTTCAGACG | | |
| PCR_Truseq_Amp_R_23 | CAAGCAGAAGACGGCATACGAGATCCACTCGTGACTGGAGTTCAGACG | | |
| PCR_Truseq_Amp_R_25 | CAAGCAGAAGACGGCATACGAGATATCAGTGTGACTGGAGTTCAGACG | | |
| PCR_Truseq_Amp_R_27 | CAAGCAGAAGACGGCATACGAGATAGGAATGTGACTGGAGTTCAGACG | | |
| pQT4MiF | ctctttccctacacgacgctcttccgatctCCATAGCATGACCGACACAT | Paired to amplify *Q* gene promoter in cv. Bobwhite | Amplifying the *Q* gene promoter to confirmaiton the sequence using Sanger sequencing. |
| pQT17MiR | ctggagttcagacgtgtgctcttccgatctCCACAGGCGTACCTGTCC | | |
| pQT17MiF | ctctttccctacacgacgctcttccgatctGTGACCACGAGCGTAAGGTT | Paired to amplify *Q* gene promoter in cv. Bobwhite | |
| pQT25MiR | ctggagttcagacgtgtgctcttccgatctAGTGGCATGGACAAAGAACC | | |
| pQT25checkF | TGCACTTGGGCAATGTAATG | Paired to amplify *Q* gene promoter in cv. Bobwhite | |
| Q5endR3 | GGAGCAGTCGTCATCATCAG | | |
| M13-F | CGCCAGGGTTTTCCCAGTCACGAC | | Sanger sequencing of *Q* gene promoter region. |
| M13R | AGCGGATAACAATTTCACACAGGA | | |
| pQT25checkF | TGCACTTGGGCAATGTAATG | | |
| pQT25checkR | AGTGGCATGGACAAAGAACC | | |
| pQT26checkF | AGGAGGCATTTAGCTTGTGGAG | | |
| pQT26checkR | GCCAGTGGGAGAAAGAGAAAGA | | |

| | | | |
|---|---|---|---|
| Q5endR3 | GGAGCAGTCGTCATCATCAG | | |
| pQT17checkF | GTGACCACGAGCGTAAGGTT | The Cas9 target flanking primers without truseq adapter tails make it eaier to amplify long fragment from wheat genomic DNA. | Long deletiondetection on Q gene promoter in M1 progeny inoculated by BSMV-pQT. |
| pQT17checkR | CCACAGGCGTACCTGTCC | | |
| pQT18checkF | AAGGAACAACGTTTCCGATG | | |
| pQT18checkR | CGCTCTCTTTCGTGTGTGTG | | |
| pQT23checkF | AATTGCACCCATCCCATTAC | | |
| pQT23checkR | AGGCCTCGTGTGTTGACC | | |
| pQT25checkF | TGCACTTGGGCAATGTAATG | | |
| pQT25checkR | AGTGGCATGGACAAAGAACC | | |
| pQT26checkF | AGGAGGCATTTAGCTTGTGGAG | | |
| pQT26checkR | GCCAGTGGGAGAAAGAGAAAGA | | |
